# Perceptual similarity representations beyond the sensory cortex

**DOI:** 10.64898/2026.10.09.757806

**Authors:** Ali Moharramipour, Zhiyong Jin, Ning Su, Yijun Ge, Takeru Matsuda, Kazuhisa Shibata, Sze Chai Kwok, Hakwan Lau

## Abstract

It is commonly thought that information supporting conscious perception is primarily represented in sensory cortices, whereas association cortex encodes more abstract, task-dependent information. However, this impression may partly stem from difficulties in experimentally assessing the functions of association areas characterized by mixed selectivity and complex spatial topography. Here, we combined repetition-suppression (RS) and multi-voxel pattern analysis (MVPA) to extract richer information from human fMRI. With this approach, we asked whether association regions represent the perceptual similarity structure among color stimuli, although this perceptual similarity information was task-irrelevant in the experiment. Our analysis revealed such representations beyond the visual cortex, in lateral prefrontal, and parietal regions. We further demonstrated an electrophysiological complement to the fMRI findings by applying the same approach to local field potentials recorded in the macaque dorsomedial parietal cortex. Our findings suggest that higher-order areas beyond the visual cortex may play a key role in subjective color perception.

## 1. Introduction

One key characteristic of neural coding underlying perception is that it preserves similarity relationships among stimuli: perceptually similar stimuli evoke similar neural representations, whereas perceptually distinct stimuli evoke dissimilar representations. Therefore, identifying neural representations that align with perceptual similarity structures can provide insight into the functional organization underlying perception (Broday-Dvir et al., 2023; Brouwer & Heeger, 2009; Contier et al., 2024; Kriegeskorte & Kievit, 2013; Kriegeskorte & Wei, 2021; Malach, 2021).

It is well documented that the structure of neural representations in the visual cortex, commonly formalized by representational dissimilarity matrices (RDMs), aligns with stimulus similarity structures (Kriegeskorte & Kievit, 2013). For example, neural RDMs from the fusiform cortex reflect face similarity structure (Chang & Tsao, 2017; Davidesco et al., 2014; Grossman et al., 2019), RDMs from V4 match color similarity structure (Brouwer & Heeger, 2009), and RDMs from ventral visual and lateral occipital cortex reflect perceived similarity structure of real-world objects (Cichy et al., 2019). In line with these findings, many hold the view that perceptual experiences are primarily shaped by representations within the sensory cortex (Block, 2005; Koch et al., 2016; Lamme, 2006; Malach, 2021).

However, it has also been argued that higher-order association areas in the prefrontal and parietal cortices are involved in subjective perception (Dehaene & Changeux, 2011; González-García et al., 2018; González-García & He, 2021; Odegaard et al., 2017; Ujhelyi et al., 2026). Specifically, it has been speculated that because human subjects seem to be able to assess how similar two perceptual stimuli are with ease, often implicitly (as in, automatically without deliberation), such information may also be available in the prefrontal cortex, regardless of task demand (Lau et al., 2022).

But the suggestion that such task-irrelevant perceptual information is represented explicitly in higher-order association areas like the prefrontal cortex is somewhat at odds with the existing literature. Numerous studies have shown that coding in the prefrontal cortex is generally shaped by task and context (Bernardi et al., 2020; Bhandari et al., 2025; Fuster, 2001; Harel et al., 2014; Mante et al., 2013; Miller, 2000; Waskom et al., 2014), with representations sometimes characterized as categorical and abstract (McKee et al., 2014; Miller et al., 2003; Pan & Sakagami, 2012; Swaminathan & Freedman, 2012).

However, the impression that task-irrelevant perceptual information may be absent in the prefrontal cortex, especially in human neuroimaging studies, may partly reflect methodological limitations. Higher-order association areas, such as the prefrontal cortex, lack the clear spatial organization characteristic of sensory cortex and contain highly mixed-selective neurons (Constantinidis & Qi, 2018; Haile et al., 2019; Ramirez-Cardenas & Viswanathan, 2016). This poses a formidable challenge for assessing the hypotheses discussed above in human subjects.

Using fMRI, we leveraged the repetition-suppression (RS) phenomenon, also known as neural adaptation, to extract RDMs at the level of single voxels. If two stimuli evoke more similar neural representations within a voxel, stronger suppression is expected when they are presented in immediate succession (Barron et al., 2016; Bhandari et al., 2019; Rigotti & Fusi, 2016). This means that the degree of RS for a stimulus pair can be taken as a measurement of how similar the two stimuli are represented by a voxel. Therefore, a corresponding RDM for a set of stimuli can be constructed when the RS effects are systematically assessed for each stimulus pair.

Conventional approaches provide an aggregate measure of each voxel’s neural population response to each stimulus, which may be insufficient for assessing representational similarity at the voxel level. In contrast, RS can directly reflect the similarity of neural coding between stimuli within a voxel. While conventional multivoxel pattern analyses (MVPA) can characterize how information is represented across a group of voxels (Haxby, 2012; Haxby et al., 2001; Nili et al., 2014), RS can provide a similar characterization, but across the neural population within a single voxel. Therefore, RS can potentially help us achieve higher effective spatial resolution when probing perceptual similarity representations (Bhandari et al., 2019; Rigotti & Fusi, 2016).

However, MVPA could still be relevant here, because the voxels whose RS-derived RDMs align with perceptual similarity (as rated by the subjects) may be sparsely distributed within an anatomical region, with their precise locations varying across individuals. Accordingly, we used MVPA methods not to assess spatial patterns of stimulus-evoked activity, but to assess *patterns of expression of the RS effects* aligned with perceptual similarity structures. In other words, we combined the strengths of both RS and MVPA.

To anticipate, using this novel method with color stimuli as the model domain, we found robust effects of perceptual similarity representations in the visual cortex, as expected. However, we also found significant effects in the lateral prefrontal and parietal cortices. These effects were observed even though perceptual similarity was irrelevant to the task being performed. The findings suggest that regions beyond the visual cortex, such as the lateral prefrontal cortex (LPFC), may be involved in visual perception.

We also conducted the same experiment in a non-human primate (NHP) to validate the approach and gain further insight into the temporal dynamics of RS effects. The NHP passively viewed the same stimuli as those presented in the human experiment, and we recorded local field potentials (LFPs) from the dorsomedial parietal cortex. The RS effects in this region reflected meaningful structure of color similarity, consistent with those identified in analogous human regions.

## 2. Results

### 2.1. Study overview

We used color stimuli to test whether representations of perceptual similarity extend beyond the sensory cortex. An RS paradigm was used to derive voxel-wise RDMs from fMRI data. This paradigm consisted of blocks presenting alternating pairs of two colors and blocks presenting single colors (Figure 1A). For each voxel, the representational dissimilarity between two colors was estimated by contrasting activity during their pair presentation with the average activity when each color was presented alone. Multiple color sets were used separately in this task (Figure 1B), yielding one RDM per color set for each voxel (e.g., Figure 2A,1). Because each color pair required a separate stimulus block, multiple smaller color sets were used instead of a single large set to maintain a manageable number of stimulus blocks while still covering a wide range of colors.

**Figure 1.**
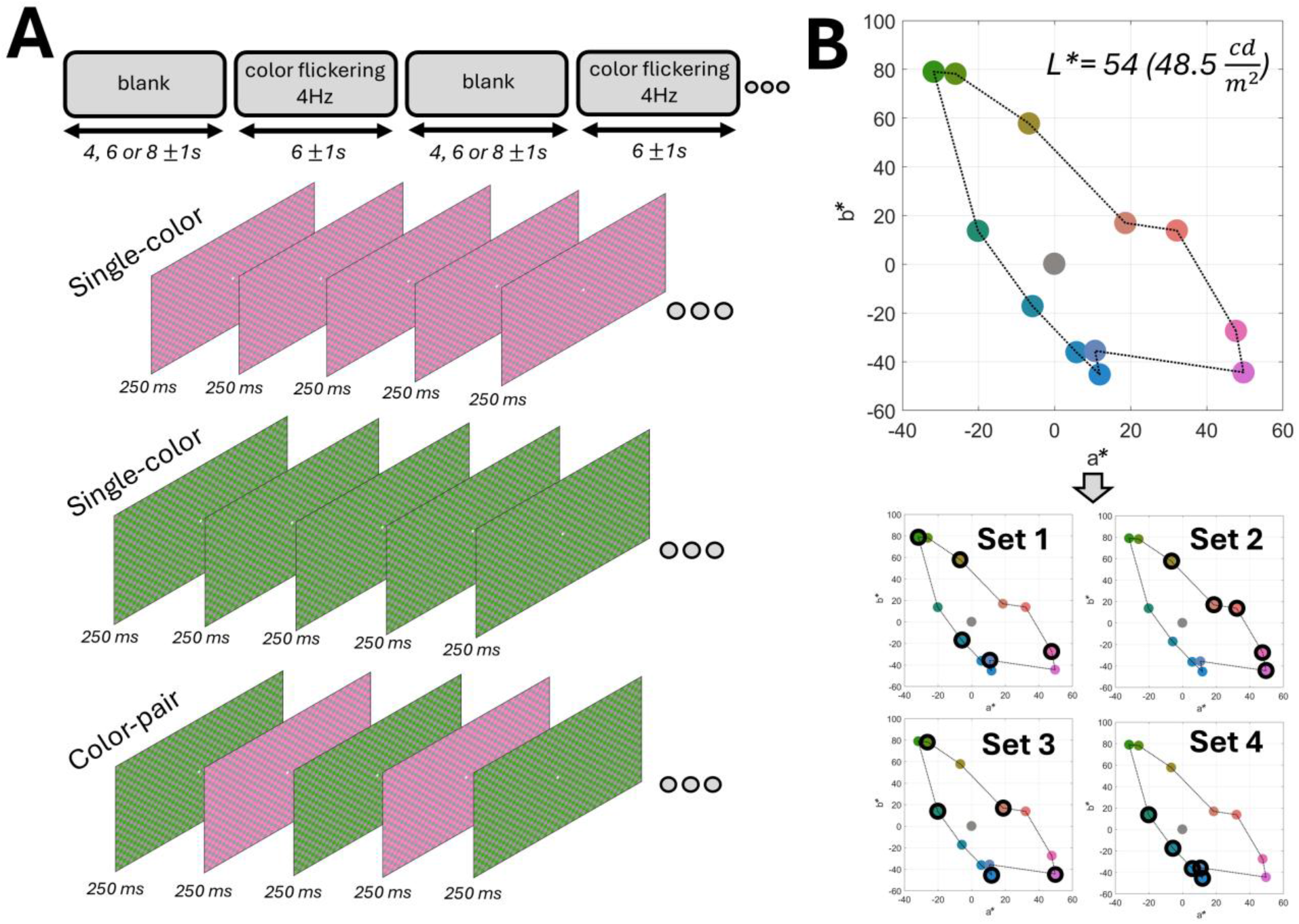
The paradigm used to extract voxel-wise neural representational dissimilarity matrices (RDMs) leveraging the repetition-suppression (RS) phenomenon. **(A)** Colors were presented as checkerboard patterns composed of a color and a default isoluminant gray in 6 ±1 s stimulus blocks, separated by blank periods. Blocks consisted of either a single color (e.g., only pink or only green) or a color pair, alternating every 250 ms (e.g., green -> pink -> green -> pink, and so on). Checkerboard elements also flickered (with gray and color elements switching) every 250 ms according to a fixed sequence. See <u>Supplementary video</u> for stimulus examples. **(B)** Isoluminant colors used in the study. Colors were divided into four sets of five colors, with each set used separately in the paradigm. The study included data from two cohorts; colors shown here were used for the second cohort (N = 4), whereas those used for the first cohort (N = 4) are shown in Supplementary Figure 2.

**Figure 2.**
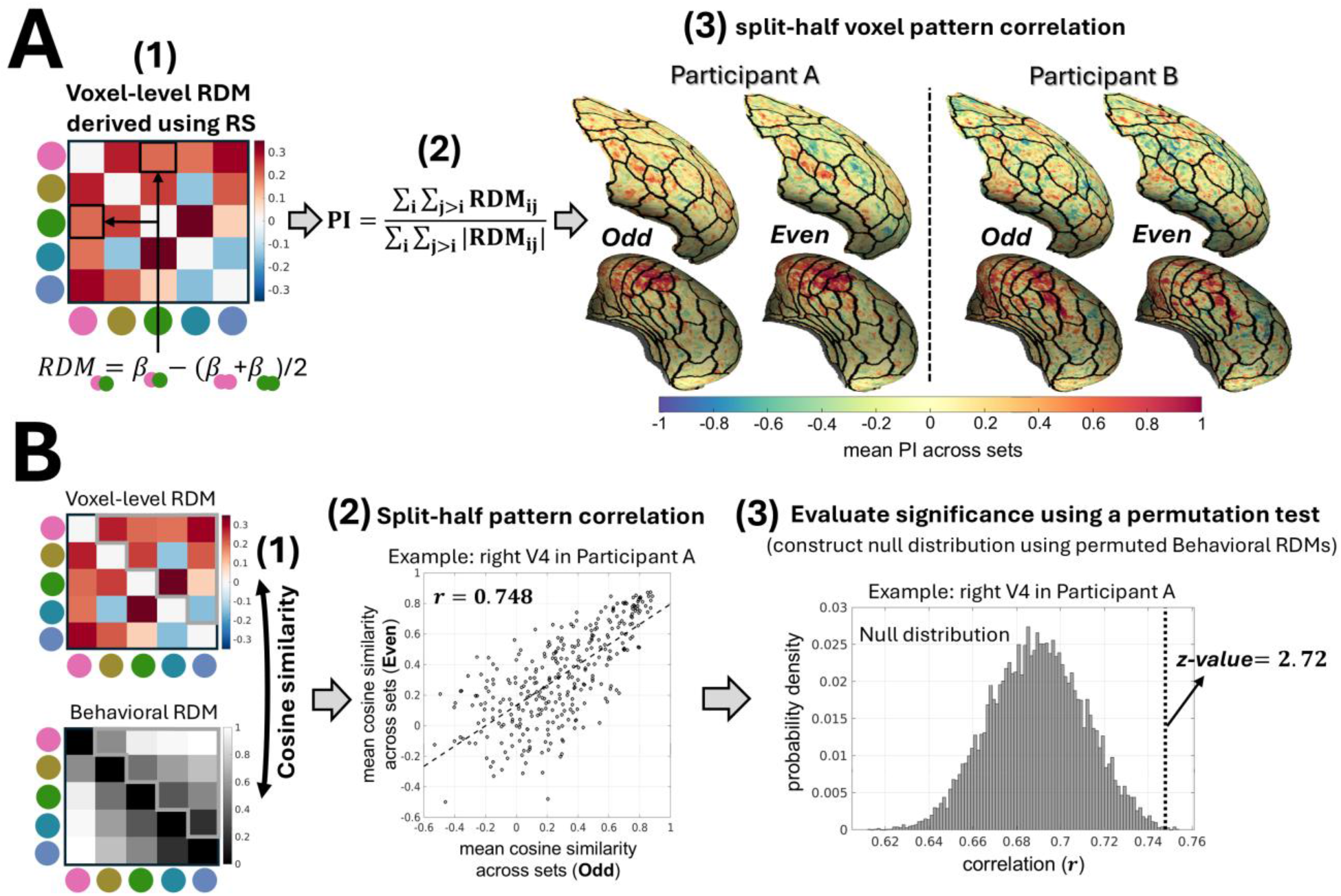
Summary of the repetition-suppression multi-voxel-pattern-analysis (RS-MVPA). **(A) (1)** For each voxel, an RDM was derived for each color set from the paradigm shown in Figure 1A, based on the RS principle. *β* represents the estimated fMRI activity associated with a given stimulus block. An example voxel RDM is shown for color set 1. **(2)** Voxels encoding color information are expected to show positive RDM cells; therefore, RDM positivity was quantified on a scale from -1 to 1 and averaged across color sets to indicate the voxel’s overall color responsiveness. **(3)** To evaluate whether an ROI contained a meaningful pattern of color-responsive voxels, data were split into even and odd runs, positivity indices were calculated separately for each half, and voxel-pattern correlations were computed between halves. The pattern correlations were transformed to z-values using Fisher’s transformation for a subsequent group-level t-test. Examples from the right lateral prefrontal cortex (LPFC) and ventral visual stream in two participants are shown. Note that examples are displayed on the average surface (fsaverage) for visualization, whereas the analyses were performed on native surfaces. **(B) (1)** Voxel-level RDMs were compared with behavioral perceptual RDMs (obtained from a separate similarity judgment task) for each color set. The upper-triangular elements were vectorized, and cosine similarity was calculated. Cosine similarity values were then averaged across color sets to reflect the overall alignment between voxel-level representation and perceptual similarity. **(2)** A split-half MVPA approach, similar to panel A, but using cosine similarity values, was applied at the ROI level. An example from right V4 in a participant is shown. **(3)** A non-parametric permutation test evaluated whether observed pattern correlations exceeded those obtained from using randomly permuted behavioral RDMs in the analysis, thereby indicating the presence of color similarity representations within the ROI. z-values were then computed and used in a subsequent group-level t-test.

A separate color-similarity judgment task (Supplementary Figure 1) was conducted for each color set to capture how each individual perceives color similarity. This task yielded one behavioral RDM for each color set for each participant (e.g., Figure 2B,1).

The study included eight participants, each scanned extensively. We conducted 36 scanning sessions and collected about 27 hours of fMRI data during the RS paradigm in total across participants. Therefore, although we had a modest number of participants, the extensive data collected per participant enabled a reasonably reliable group-level analysis.

In this Results section, we first identify candidate regions containing color-responsive voxels. Next, by comparing neural RDMs with behavioral RDMs, we assess which regions contain representations of perceptual similarity. Lastly, we present results from LFP recordings in an NHP that was presented with the same RS paradigm as the human participants.

### 2.2. Candidate regions with color-related responses

Representational dissimilarity (i.e., each cell in voxel-level neural RDMs) can, by definition, take on negative values (Figure 2A,1). However, for voxels that represent color, the values are expected to be positive, given the RS principle. This is because responses to alternating pairs of colors should lead to less suppression (i.e., higher activity) than responses to colors presented individually. We therefore calculated a positivity index for each voxel, ranging from -1 to 1 (Figure 2A,2), to quantify color-relevant responses. A value of 1 indicates that all RDM cells are positive and the voxel is likely color-responsive. However, due to fMRI noise, the positivity index may not reach 1, but is expected to be above zero for color-responsive voxels.

We then used an MVPA-based approach (Figure 2A,3) to identify ROIs containing reliable patterns of color-responsive voxels, with color responsiveness quantified by the voxel positivity index. This ROI-level analysis increases sensitivity by accounting for the possibility that responsive voxels may be distributed rather than forming clear clusters across participants within a region. This consideration is particularly relevant in higher-order areas; for example, compare the LPFC and ventral visual stream in Figure 2A,3.

**Figure 3.**
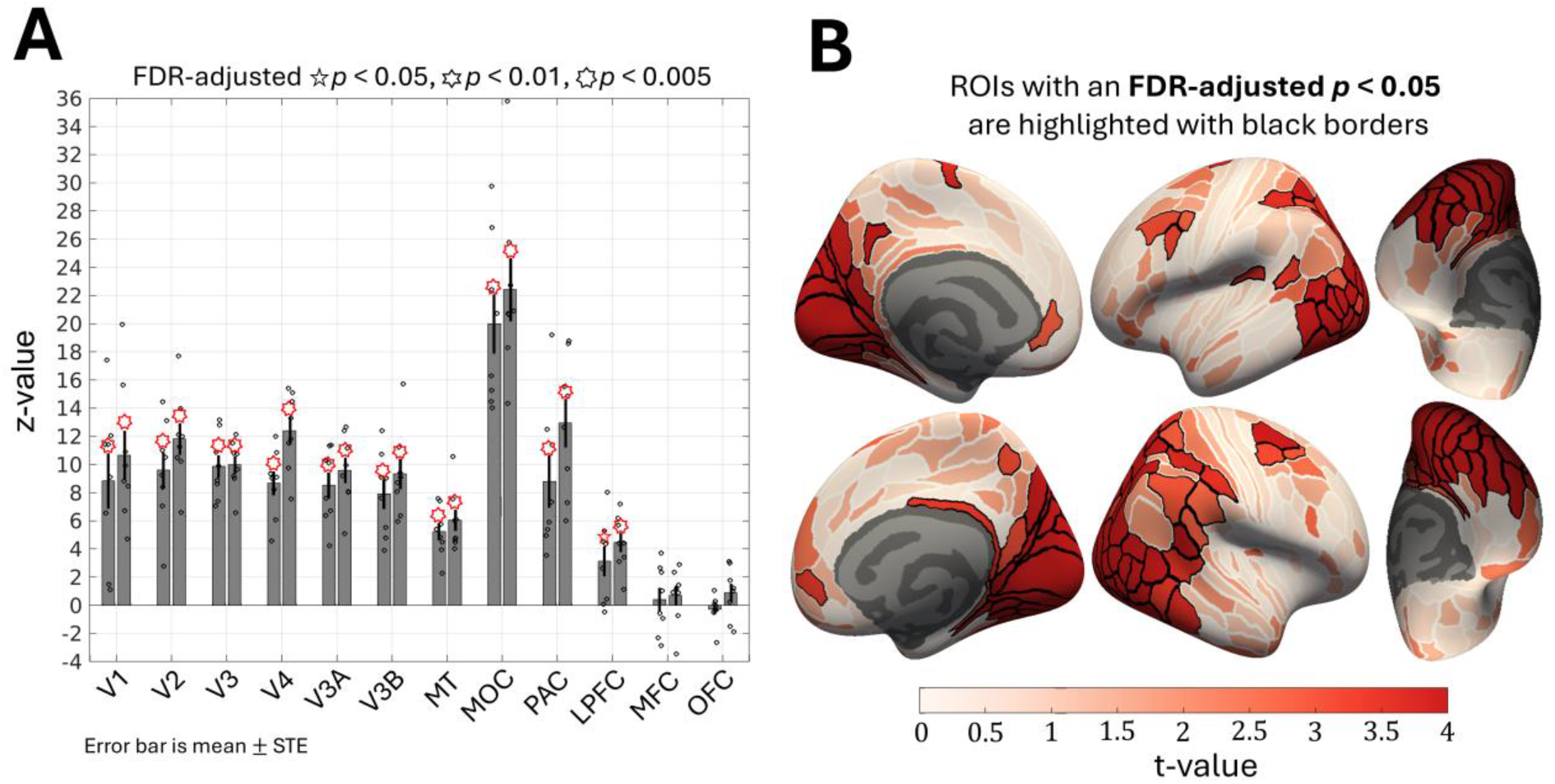
RS-MVPA results identifying candidate ROIs with consistent color-related responses (see Figure 2A for the analysis summary). **(A)** Bars show mean participant z-value (±STE), with dots indicating individual participants. Left and right bars correspond to the left and right hemispheres, respectively. Visual areas V1 to MT were delineated from individual retinotopic maps (Supplementary Figure 3). LPFC, parietal association cortex (PAC), medial frontal cortex (MFC), and orbitofrontal cortex (OFC) were defined by grouping parcels from Glasser’s atlas (Glasser et al., 2016) based on their anatomical locations (Supplementary Figure 3B). MOC was manually defined to encompass medial occipital regions not covered by retinotopically delineated ROIs, including regions around the parieto-occipital sulcus (POS) (Supplementary Figure 3). All visual areas, PAC, and LPFC showed significant color-related responses, as indicated by a group-level t-test on individual-level z-values (FDR-adjusted *p* < 0.05). **(B)** Results on the fine-grained Glasser’s atlas (Glasser et al., 2016). The color bar represents group-level t-values, and significant ROIs (FDR-adjusted *p* < 0.05) are highlighted with black borders.

We pre-registered a set of ROIs (supplementary Figure 3B), and the RS-MVPA results for these ROIs are shown in Figure 3A. Not only visual areas, including V1, V2, V3, V4, V3A, V3B, MT, and the medial occipital cortex (MOC), but also higher-order areas, including the parietal association cortex (PAC) and LPFC, showed consistent patterns of color-related responses. Note that the reported z-values were computed parametrically using Fisher’s z-transformation and scaled according to the number of voxels, as required for proper statistical inference; thus, larger ROIs (e.g., MOC is about seven times larger than V4) yield higher z-values for the same level of response pattern consistency. However, spatial smoothness may inflate these z-values as nearby voxels are not entirely independent. Nevertheless, we assessed significance at the group level using a t-test on individual-level z-values. This second-level statistic, by relying on inter-participant variance, helps protect against potential biases related to spatial smoothness. Furthermore, because the number of voxels scales these individual-level z-values, comparisons across ROIs should likewise be based on the group-level measure. Under this group-level statistic and after the FDR correction (adjusted *p* < 0.05), significant effects were observed in the PAC and LPFC, in addition to visual areas.

To extend our analysis beyond the predefined ROIs, we applied RS-MVPA to fine-grained Glasser’s atlas (Glasser et al., 2016). As shown in Figure 3B, beyond visual areas, several association regions showed consistent patterns of color-responsive voxels, including the PAC, particularly around the intraparietal sulcus, the posterior parts of the LPFC, and areas near the temporo-parietal junction.

For comparison, we also generated a simple functional color localizer map by contrasting colored versus grayscale naturalistic images presented in our retinotopic paradigm (Supplementary Figure 4). Qualitatively, this localizer showed strongest activity in V4 and more anterior ventral visual areas, with subtle yet noticeable activity in some association regions consistent with those identified by the RS-MVPA analysis. However, some ROIs detected by RS-MVPA showed no readily visible color-related activity in the localizer. This could reflect the sensitivity of the RS-based approach to complex neural responses: even if color is encoded, responses to colored and grayscale images may appear similar enough at the aggregate neural population level within a voxel that a simple activation contrast may not detect this information.

That said, this comparison may not be entirely fair because the RS-MVPA analysis was conducted at the ROI level and used substantially more data than the localizer. Furthermore, RS-MVPA may not be entirely free from false positives. We cannot dismiss the possibility that non-color-related factors could also influence the positivity index; for example, stronger responses to the presence of two stimuli than to one, regardless of color identity (e.g., reflecting stimulus counting), could result in a high positivity index value. Such factors could inflate pattern consistency and lead to false positives. We therefore treat the results here as candidate regions with color-related responses and use a more stringent analysis in the next section to identify regions containing perceptually relevant color representations rather than a general color response.

### 2.3. Regions with representations of perceptual color similarity

Voxel-level neural RDMs were first compared with behavioral RDMs using cosine similarity (Figure 2B,1). We then assessed whether the pattern of these voxel-wise cosine similarity values was consistent within each ROI (Figure 2B,2). Crucially, to determine whether this consistency was genuinely driven by perceptually structured color information, we compared the observed value against a null baseline generated from using randomly permuting behavioral RDMs in the analysis (Figure 2B,3), yielding a z-value for each ROI. Significance was then assessed at the group level using a t-test on individual-level z-values, followed by FDR correction across ROIs.

This analysis revealed representations of perceptual color similarity in most visual areas, strongest in V4 and MOC and weakest in V1 and V3A, as indicated by the z-values in Figure 4A. Beyond visual areas, higher-order regions, including PAC and LPFC, also showed significant effects. Unlike the z-values in Figure 3A, the z-values here were obtained from a stringent nonparametric test and are therefore unlikely to reach as high values. For example, based on a simulation approximating the actual experimental conditions and design, a z-value of about 3.5 is considered exceptionally high (Supplementary Figure 5).

**Figure 4.**
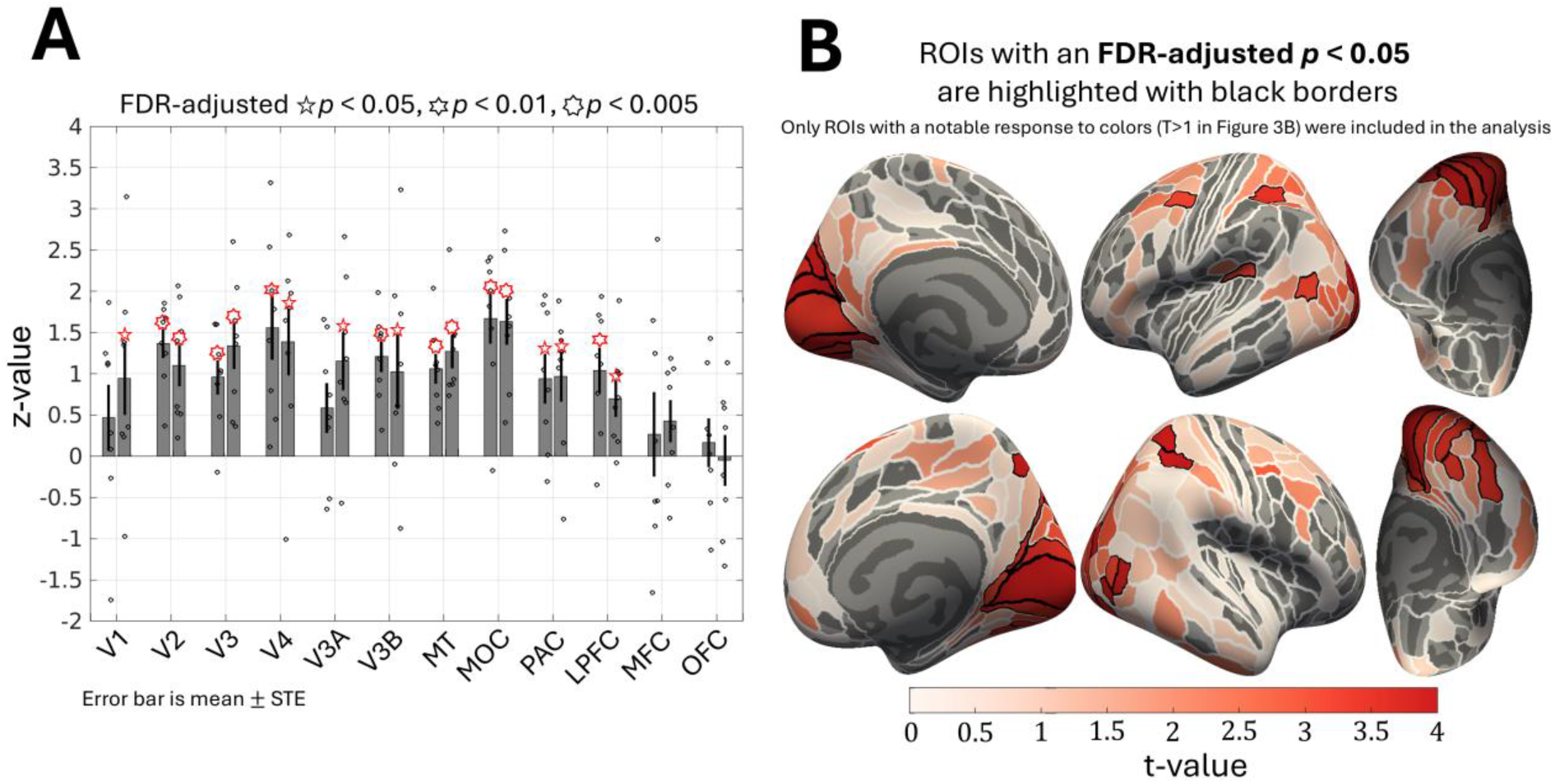
RS-MVPA results identifying ROIs with representations of perceptual color similarity (see Figure 2B for the analysis summary). **(A)** Bars show mean participant z-values (±STE), with dots indicating individual participants. Left and right bars represent left and right hemispheres, respectively. Most visual areas, PAC, and LPFC showed significant effects, as indicated by a group-level t-test on participant z-values (FDR-adjusted *p* < 0.05). **(B)** Fine-grained Glasser’s atlas (Glasser et al., 2016) results. Only ROIs with notable color-related responses in the first analysis stage (t-value > 1 in Figure 3B) were analyzed to alleviate the multiple-comparison burden. The color bar represents group-level t-values and significant ROIs (FDR-adjusted *p* < 0.05) are outlined with black borders. Beyond visual areas, apparent effects were observed around the intraparietal sulcus in PAC and in the posterior LPFC. See Supplementary Figure 6 for results from alternative group-level tests that showed additional ROIs within PAC and LPFC surviving FDR correction.

Figure 4B shows the results from the fine-grained Glasser’s atlas (Glasser et al., 2016). As in Figure 4A, most visual areas exhibited representations of perceptual color similarity. Within MOC, this included V6, DVT, and Prostriata. Beyond visual areas, a few ROIs in the PAC and LPFC also showed perceptually relevant representations of color similarity. However, it is possible that some ROIs with meaningful effects may not have emerged as significant, given the large number of ROIs tested and the modest number of participants used in the t-test. We therefore also applied complementary group-level tests that may be more sensitive for modest sample sizes, including Fisher’s combined probability test (Fisher, 1970), Stouffer’s test (Whitlock, 2005), and a nonparametric bootstrap test (Mooney et al., 1993). Using each of these tests, more ROIs survived FDR correction, notably in the posterior LPFC (55b, PEF, 8Av, 8C, 6a) and around the intraparietal sulcus in the PAC (AIP, 7PC, MIP, LIPv, IPS1, IP0, IP1, IP2, VIP, PFt) in at least one hemisphere (Supplementary Figure 6).

In summary, the results of this section suggest that representations of perceptual similarity are not limited to the visual cortex, but extend to higher-order regions, including the LPFC, particularly its posterior areas, and the PAC, especially regions around the intraparietal sulcus.

#### 2.3.1. Exploratory observations

We conducted several exploratory analyses to characterize the granularity of perceptual similarity representations across regions, their connectivity profile, and their functional relevance during performing the similarity judgment task, as described below.

We used two sets of closely spaced colors (near sets; sets 2 and 4 in Figure 1B) and two sets of more widely spaced colors (far sets; sets 1 and 3 in Figure 1B). Representations of perceptual similarity in the near sets would likely require finer-grained color tuning, whereas coarse-grained tuning could be sufficient for the far sets. Specifically, distinguishing among different orange/pinkish or bluish colors would likely be necessary to effectively represent perceptual similarity within the near sets, whereas a coarser representation that does not distinguish, for example, among shades of blue could suffice for the far sets. We therefore tested whether ROIs contained voxels with finer- or coarser-grained representations, aligned with perceptual similarity. We contrasted the ROI-level consistency measures (same as Figure 2B,2) obtained separately from near and far sets, with significance assessed using a nonparametric permutation approach (similar to Figure 2B,3; using permuted RDMs in computing this contrast). Most visual areas showed a trend toward finer-grained color representations, which was most pronounced in V3 and V4. A similar trend, though not as pronounced as in V3 and V4, was also observed in higher-order areas, including the LPFC (Supplementary Figure 7).

We examined the whole-brain functional connectivity of voxels with representations of color similarity using fMRI data collected while participants watched the first 15 or 20 min of the movie *Interstellar*. Movie watching provides naturalistic stimulation and allows functional connectivity to be assessed without an explicit behavioral task. These connectivity maps provided additional insight. For example, the relevant color-similarity voxels in V4, an area well known for color processing, exhibited apparent functional connectivity with areas within MOC, including regions around the POS, particularly DVT, as well as some areas within the PAC (around the intraparietal sulcus), and posterior LPFC (Supplementary Figure 8).

The RS paradigm involved no task related to color similarity. However, we were able to explore brain activity related to actively using such similarity information from fMRI data collected while participants performed the color-similarity judgment task. This analysis qualitatively showed that some regions with representations of perceptual similarity, in both visual and non-visual areas, may also be engaged during the active task of judging color similarity, with somewhat stronger engagement in non-visual areas (Supplementary Figure 9).

Finally, we note that we also conducted a pre-registered ‘univariate’ analysis of RS-fMRI data. In this analysis, a design matrix was specifically constructed to enable one of its regressors to capture voxels whose RS profiles were aligned with the behavioral RDMs. However, we subsequently found it to be inefficient and potentially problematic because of considerable multicollinearity among the regressors. The results and methodological details are therefore presented only in Supplementary Figures 10 and 11. Despite these limitations, this analysis suggested that MOC and V4 contained the highest proportions of voxels with representations of perceptual color similarity, with a progressive increase from V1 to V4. Such voxels were also observed within PAC and LPFC.

### 2.4. Non-human primate electrophysiology

We conducted the same experiment as in Figure 1A on an NHP, with 32 microelectrodes implanted in the left dorsomedial parietal cortex, spanning V2, V6A, MIP, and PEc (Figure 6). We obtained extensive LFP recordings during stimulus presentation. This allowed us to more directly examine RS effects with high temporal resolution and test whether V2, V6A (situated near the POS), and MIP (located in the medial bank of the intraparietal sulcus) show representations of color similarity consistent with the human fMRI findings (Figure 4B; MIP survived as significant in Supplementary Figure 6). In the human experiment, participants performed a task to remain engaged (see section 4.1.2.1), but the task did not require them to compare the similarity between colors in any way. Likewise, in the NHP experiment, the monkey passively viewed the stimuli while simply maintaining fixation, without performing any task related to color similarity.

LFP power increased during stimulus presentation, with a dominant gamma-band component at 30-80 Hz (see the example electrode in Figure 5A). The gamma band is commonly thought to reflect engagement of local neural populations and synchronized synaptic activity (Jia & Kohn, 2011; Ray & Maunsell, 2010). We used the gamma-band spectral centroid, the frequency at which power is concentrated, as a proxy for local population engagement. Increases in the spectral centroid could reflect an elevated local excitatory drive (Ray & Maunsell, 2010). As shown in the exemplar electrode from area V2 (Figure 5B), the spectral centroid decreased toward a relatively steady state. It was higher in color-pair blocks than in single-color blocks throughout stimulus presentation from the first color alternation onward (250 ms onward), with the difference being particularly large during the steady period. These patterns are highly consistent with the expected effects of RS.

**Figure 5.**
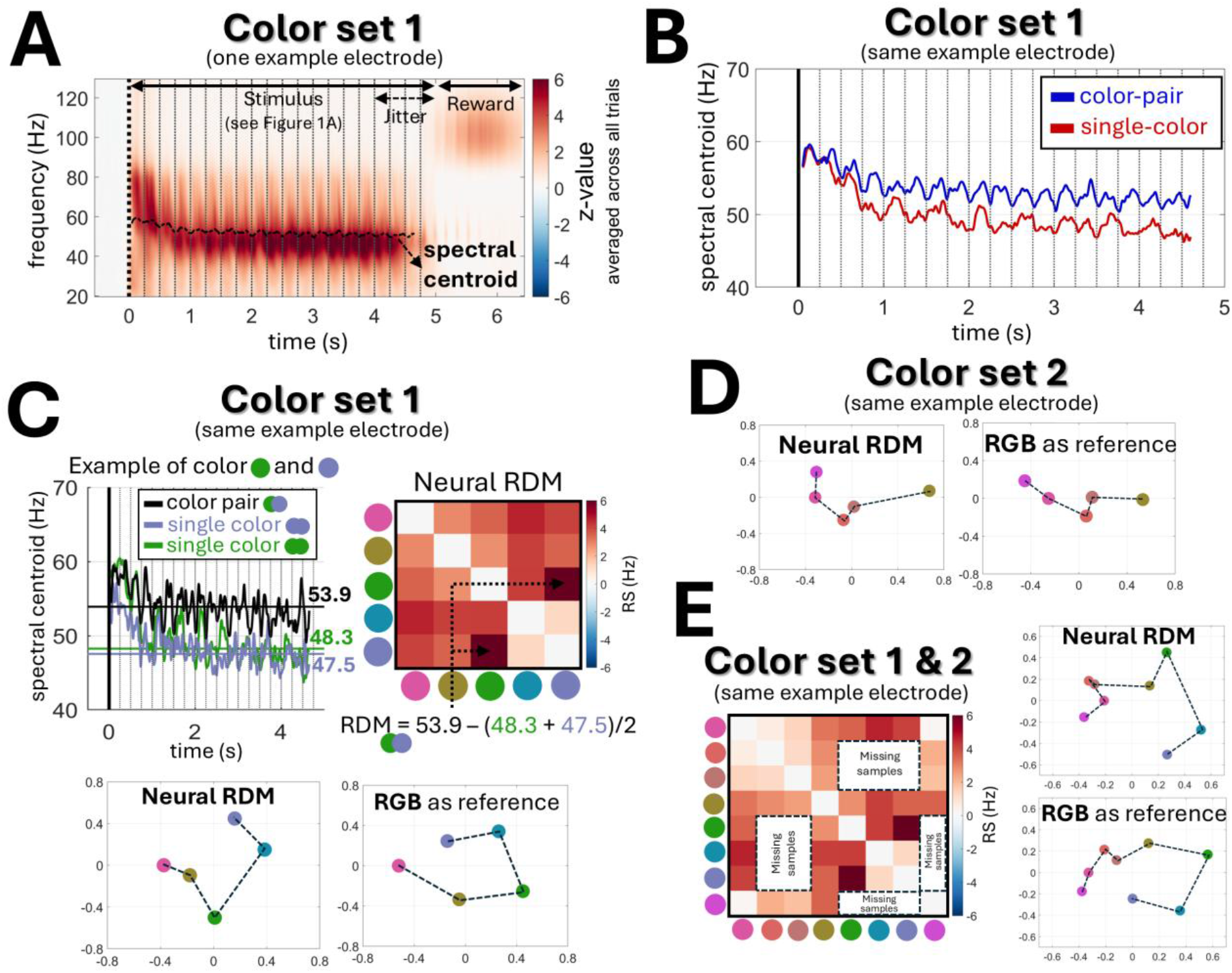
RS-based analysis of local field potentials (LFPs) recorded in a non-human primate (NHP). The same stimuli as Figure 1A were presented for 4.5 ± 0.5 s, and the NHP received a reward for maintaining central fixation. Color sets 1 and 2 (Figure 1B) were used in the experiment. Results are shown for a single exemplar electrode in area V2. **(A)** This panel shows the time-frequency response, z-normalized relative to baseline and then averaged across all trials from color set 1. The spectral centroid, reflecting the weighted-average frequency at each time point, was then computed in the gamma band (30-80 Hz). Time 0 s indicates the stimulus-block (trial) onset, and dotted lines denote color-flickering onsets (every 250 ms). **(B)** This panel shows the spectral centroid computed separately from all color-pair trials and all single-color trials corresponding to color set 1. **(C)** Example spectral centroid responses for a color-pair (green-violet) and the corresponding single-color trials (green or violet only) are shown in the top-left plot. The median spectral centroid during the steady period (about 1.5 s after stimulus onset) was used as a summary response measure to then construct RDMs analogous to Figure 2A,1. The resulting RDM for the exemplar electrode from color set 1 was visualized in 2D space alongside a reference representation based on RGB color values. **(D)** The same procedure was applied to color set 2 trials for the same electrode, with the resulting RDM visualized in a 2D space alongside a reference RGB-based representation (see Supplementary Figure 12 for the panels A-C corresponding to color set 2). **(E)** RDMs from color sets 1 and 2 (panels C and D) were combined and visualized. Two colors were shared across sets, resulting in a combined 8 × 8 matrix. Some RDM cells are missing, as the experiment was run separately for each color set; only available cells were used to generate the 2D visualization.

We used the spectral centroid responses from each of the color-pair and single-color conditions to construct neural RDMs following the same framework as the fMRI analysis (Figure 5C). For the exemplar V2 electrode, these RDMs closely matched expected color similarity structures (Figures 5C-E). This suggests that gamma-band spectral centroids can sensitively capture RS effects elicited by our experimental paradigm, which can then be leveraged to extract underlying neural representations.

Across electrodes, neural RDMs showed high positivity indices (Figure 6B), indicating robust RS effects at nearly all sites. Neural RDMs also showed significant alignment with expected color similarity structures in V2, V6A, and MIP (Figure 6C). Although PEc coverage was very limited, electrodes clearly situated within PEc showed no significant effect, providing only weak evidence for representations of color similarity in this region. These findings appear consistent with those from our human fMRI study (Figure 6D).

**Figure 6.**
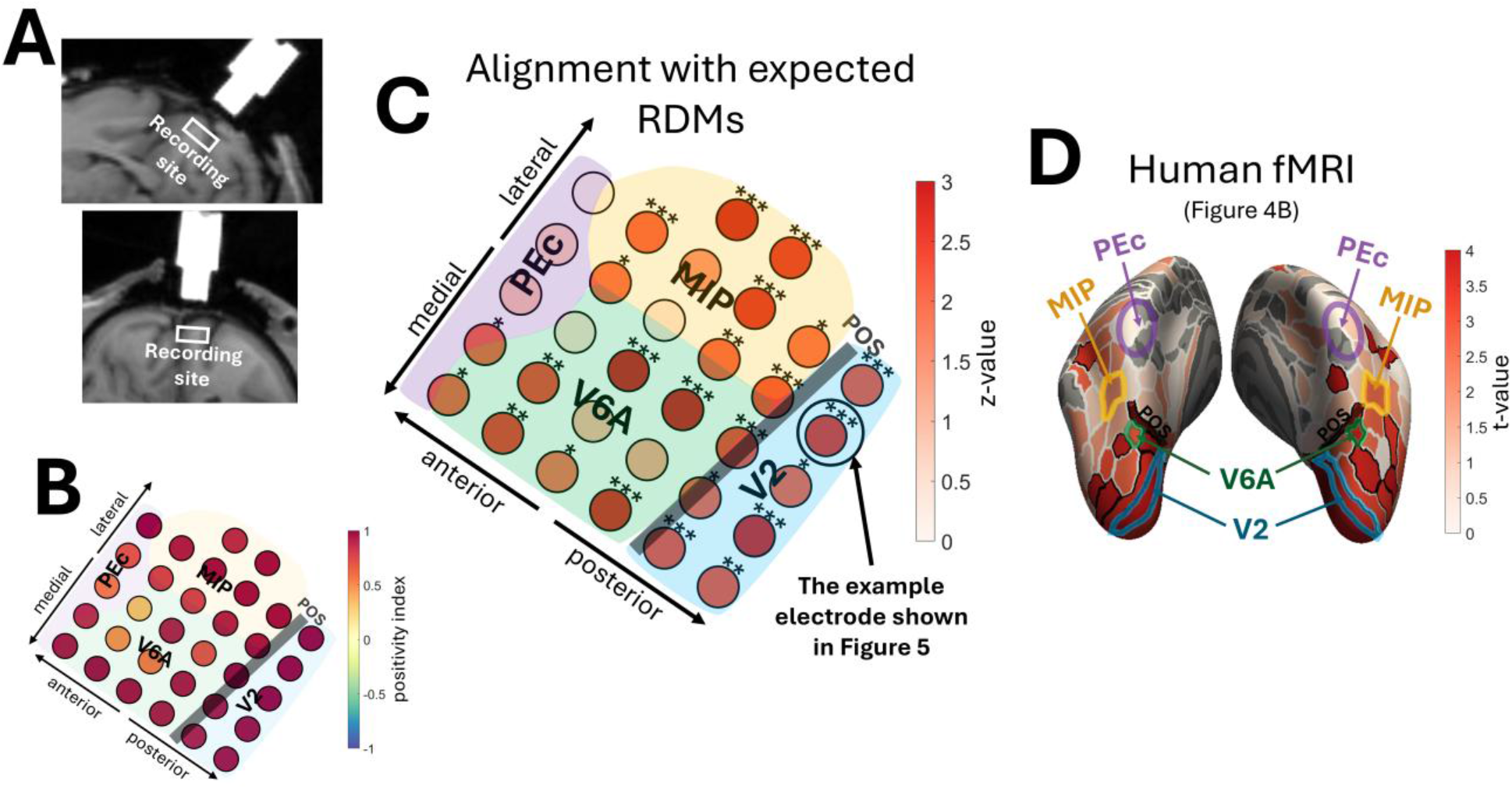
Alignment between RS-derived neural RDMs and expected RDMs across individual NHP electrodes. **(A)** Structural MRI slices showing the electrode chamber and recording site. Most electrodes were located anterior to the POS in V6A, MIP, and PEc, while some were positioned posterior to the POS in V2, according to the D99.v2.0 macaque atlas (Saleem et al., 2021). **(B)** Positivity indices (Figure 2A,2) of neural RDMs are shown for each electrode. Most electrodes showed high positivity indices, suggesting the presence of color-relevant information. **(C)** Neural RDMs were compared with RGB-based reference RDMs using cosine similarity (Figure 2B,1), with significance assessed using a nonparametric (Mantel permutation) test (FDR-adjusted \**p*<0.05, \*\**p*<0.01, \*\*\**p*<0.005). The color bar indicates the test z-values. Results were highly similar when L*a*b*-based reference RDMs were used instead. The electrodes near or between two anatomical landmarks were visualized as crossing both landmarks, reflecting uncertainty in their anatomical assignment. **(D)** Human fMRI RS-MVPA result (same as Figure 4B) with V2, V6A, and MIP marked according to Glasser’s atlas (Glasser et al., 2016). Because PEc has no explicit homolog in this atlas, its approximate location is marked based on a previous study (Pitzalis et al., 2019). While these areas are anatomically adjacent in the NHP brain, they are spatially separated in the human brain according to standard atlases. Consistently, both NHP electrophysiology and human fMRI showed evidence for perceptually relevant color representations in V2, V6A, and MIP.

Notably, we applied the same RS-MVPA approach used in our fMRI analysis (Figure 2) to this 32-channel electrode array, treating each electrode as a voxel. The analysis revealed significant color-related responses (z-value = 4.4, *p* < 0.001; Figure 2A analysis) and representations of color similarity (z-value = 2.04, *p* = 0.005; Figure 2B analysis), consistent with the results shown in Figures 6B and 6C, respectively.

We used the median spectral centroid values during the steady period (1.5 s after stimulus onset) to derive neural RDMs (Figure 5C). However, analysis at each time point (Supplementary Figure 13) showed that RDMs reflecting color similarity could already be derived following the first color alternation (250-500 ms from stimulus onset). Essentially, the RS induced by presenting color 1 and then color 2, each for 250 ms, was already sufficient to measure representational dissimilarity between colors 1 and 2, without requiring further alternations. However, additional alternations and taking the median of their responses reduced noise, allowing for a more robust estimation of the RDMs.

Gamma-band LFPs are highly localized, with a spatial spread as small as 400 μm (Ray & Maunsell, 2010). However, to minimize the risk that our findings are confounded by volume conduction, we repeated the analysis shown in Figure 6 using bipolar referencing. Results remained consistent, with areas V2, V6A, and MIP showing substantial evidence for representations of color similarity (Supplementary Figure 14). As a side note, the high-gamma power observed during the reward period (e.g., Figure 5A) disappeared entirely with bipolar referencing across all electrodes, suggesting that this signal had a non-neural origin, likely reflecting EMG contamination of the dural reference from jaw or tongue movements during reward.

Lastly, to demonstrate the advantage of the RS-based approach, we compared it with a non-RS method based on evoked responses to individual colors. These responses were recorded in a control experiment in which the NHP viewed each color individually, presented as a solid square. Although this approach captured some color structure, it provided only a low-resolution representation at each electrode that failed to show statistically significant alignment with the expected representation of color similarity (Supplementary Figure 15). Essentially, although aggregate neural population responses, as reflected in the response of an electrode, can show stimulus selectivity (i.e., higher responses to certain stimuli), such responses may provide only limited information about the structure of stimulus representations across the neural population. In contrast, the RS approach could recover this representational structure in detail.

## 3. Discussion

Using color stimuli, the present study employed an fMRI RS-MVPA approach and found that representations of perceptual similarity are present not only in visual areas but also in higher-order regions, including the PAC (particularly around the intraparietal sulcus) and the LPFC (particularly its posterior portion) (Figure 4). Notably, perceptual similarity relationships among stimuli were not relevant to the task being performed, yet these higher-order regions nonetheless represented such relationships, which may highlight a possible contribution of these regions to everyday perception.

Direct LFP recordings from an NHP dorsomedial parietal cortex during passive viewing revealed similarity representations consistent with the human fMRI findings (Figure 6). This LFP analysis provided direct insight into the neural dynamics of RS and its modulation by stimulus similarity. Specifically, it demonstrated that shifts in spectral centroid within the gamma band can serve as a sensitive measure of RS effects reflecting representational similarity, a feature that, to our knowledge, has not been previously reported (Figure 5).

### 3.1. Higher-order association areas

Previous fMRI studies using RS have shown similarity representations in visual areas, but evidence from higher-order association regions has been limited, with reported effects mainly involving categorical and conceptual representations (Aguirre, 2007; Bird et al., 2014; Gotts et al., 2011; Persichetti et al., 2015). These studies typically modeled RS-fMRI responses parametrically as a function of stimulus similarity or binary category membership, rather than extracting voxel RDMs and comparing them with behaviorally derived perceptual RDMs. By incorporating an MVPA approach, we further accounted for the possibility that relevant voxels may be distributed rather than organized into spatially consistent clusters, particularly in association areas, such as the LPFC, given their multifunctional roles and non-uniform spatial organization (Constantinidis & Qi, 2018; Ramirez-Cardenas & Viswanathan, 2016). This approach, therefore, enabled us to more sensitively probe these regions than conventional RS-based analyses and revealed representations of perceptual similarity in the LPFC and PAC.

Consistent with our findings, previous studies in macaques have identified visual representations in the LPFC even in the absence of task demands (Bellet et al., 2022; Chan, 2013; Haile et al., 2019; Schwedhelm et al., 2020), including responses to color (Haile et al., 2019; Schwedhelm et al., 2020). Human studies have also shown that visual features can be decoded from LPFC fMRI activity (Lin & Lau, 2024) and have reported visual responses in this region, such as to faces (Chan, 2013; Rajimehr et al., 2009). However, LPFC representations have often been regarded as abstract, categorical, and semantic-like (Miller et al., 2003; Pan & Sakagami, 2012; Shoham et al., 2025). Our findings may extend this view by suggesting that LPFC representations may not be limited to coarse categorical distinctions but also preserve graded perceptual similarity structures, even among closely related colors (Supplementary Figure 7).

The PAC, particularly around the intraparietal sulcus, also showed representations of perceptual similarity. Comparable representations were also observed in macaque MIP, which is located in the medial bank of the intraparietal sulcus. These intraparietal regions, including MIP, are known to exhibit visual sensory responses (Arcaro et al., 2011; Seelke et al., 2012). Beyond general visual responses, and relevant to our observation, these areas have been reported to carry visual stimulus-identity information, for example, for stimuli defined by conjunctions of color and motion-direction (Ibos & Freedman, 2017), for face or object identity (Jeong & Xu, 2016), for semantic object similarity and object identity, even under passive viewing (Neyens et al., 2017), and mixed selectivity for various categories of ethogram features (Wang et al., 2024). We used anatomical atlases to establish a correspondence between the human and macaque regions. At the broader anatomical level, the recorded region in the macaque largely encompassed the dorsomedial parietal cortex. Our observed representations of similarity in this area may be in line with the higher-level sensory and cognitive information previously reported in neurons within this region (Wang et al., 2024; Zhou et al., 2026).

### 3.2. Visual areas

Among retinotopic visual areas, V4 showed the strongest effect for representations of perceptual color similarity, while V1 showed the weakest (Figure 4A). V3 and V4 also exhibited finer-grained representations than the other areas (Supplementary Figure 7). These findings are consistent with previous studies, identifying V4 as a key region for color processing and suggesting increasing specialization of color representations from V1 to V4 (Brouwer & Heeger, 2009; Conway et al., 2007; Kim et al., 2020).

We also observed representations of color similarity in area MT, V3A, and V3B. Although these regions are not commonly considered color-processing areas, previous studies have reported color-related responses, albeit of small magnitude, in MT (Conway, 2014) and V3A (Tootell et al., 2004; Tootell & Nasr, 2017), and implicated V3B in color filling-in (Gerardin et al., 2018). A previous fMRI MVPA study also showed above-chance color decoding in MT and V3AB, although the decoding was substantially weaker than in V1-V4, and did not reflect perceptual similarity (Brouwer & Heeger, 2009). In contrast, our results suggest that color representations in these areas are aligned with perceptual similarity. Our RS-MVPA approach may detect complex or spatially sparse representations that conventional MVPA can miss, particularly when information is represented locally within a few voxels, rather than being distributed across voxels within a region. This may explain the discrepancy with the previous fMRI study, as MT and V3AB regions are likely not strongly specialized for color processing in the same way as ventral visual regions such as V4 are (Conway, 2014).

We further observed a strong effect reflecting representations of color similarity in area MOC, encompassing regions associated with the visual periphery, as well as areas around the POS, including DVT, V6, and partly V6A; DVT, V6, and V6A also showed the effect when analyzed individually (Figure 4B). This finding was unexpected in our initial analyses, although we pre-registered for this ROI after observing an effect within this area in our first cohort. Also, previous studies that employed an fMRI RS-based approach reported comparable findings regarding color in the MOC (Aguirre, 2007; Bird et al., 2014).

While color coding has been reported in regions associated with peripheral vision (Aguirre, 2007; Conway et al., 2007; Lafer-Sousa & Conway, 2013), DVT, V6, and V6A are not traditionally recognized as processing color information. These areas are typically associated with visual motion, self-motion, and visuomotor processing (Galletti & Fattori, 2018; Pitzalis et al., 2010). However, to our knowledge, color-related signals have been reported in V6 and V6A, albeit weakly in a prior fMRI study (Pitzalis et al., 2018). Moreover, we observed functional connectivity between V4, a well-established color-processing region, and DVT (and, to a lesser extent, V6 and V6A), which may provide additional support for our findings (Supplementary Figure 8).

Critically, we observed a similar finding in V6A using the same RS-based approach with macaque LFP recordings (Figure 6). Together with our fMRI results, our study provides novel evidence that V6A contains color representations organized in a perceptually meaningful way. Previous studies in macaques have shown that V6A encodes visual object attributes, such as shape and orientation, even during passive viewing without motor action (Breveglieri et al., 2015; Fattori et al., 2012), as well as action-relevant properties such as object affordance (graspability) (Breveglieri et al., 2015). It is therefore conceivable that color information may also contribute in this context, for example, by interacting with shape information to guide actions such as grasping ripe fruits.

These findings, however, do not necessarily imply specialized color selectivity, in the sense that neurons show consistent responses to a hue largely independent of brightness, as reported in ventral visual areas such as V4 (Conway et al., 2007). Although we used isoluminant colors, the observed effects could arise from multiselective neural populations that are not strictly color-selective, but rather respond to color as well as other visual features, such as luminance, motion, or spatial structure. It is noteworthy that our observation that dorsal regions (MT, V3AB, DVT, V6, and V6A) showed representations of color similarity, in addition to the ventral regions, aligns with emerging evidence that the dorsal stream also represents perceptually relevant visual information, even when no visuomotor response is required (Freud et al., 2020).

### 3.3. Potential limitations

Previous studies have cautioned that RS may be inherited from upstream regions, and therefore may not always reflect stimulus-selective responses (Larsson et al., 2016; Larsson & Harrison, 2015; Mur et al., 2010). Downstream neurons invariant to specific stimulus information could still exhibit RS if they receive projections from earlier neurons showing RS, thereby leading to false interpretations of stimulus selectivity. However, our study, beyond identifying the presence of RS (Figures 3 and 6B), importantly tested whether RS magnitude matched specific RDMs (Figures 4 and 6C). A false-positive effect in this analysis would require the graded similarity structure, rather than simply the presence of RS, to be inherited from upstream areas. This scenario is probably less likely because neural responses are typically nonlinear with respect to their inputs; therefore, similarity structure may fade in invariant neurons (especially in higher-order areas where information passes through several processing stages), while the general RS effect could still remain. Empirically, there were regions or electrodes that showed a marked effect for the presence of RS (Figures 3B and 6B), but almost no effect of alignment with perceptual similarity (Figures 4B and 6C). Accordingly, our secondary analysis regarding perceptual similarity is more specific and may also be less prone to inheritance effects, although this confound cannot be entirely ruled out.

Certain low-level stimulus factors, such as chromatic aberration (e.g., color fringing near checkerboard edges) and potential mismatches between subjective and physical isoluminance, could in principle influence neural signals. These effects, however, are expected to mostly influence early visual areas. Importantly, our primary focus was on higher-order association areas, where these low-level factors are less likely to be a major concern; even so, they are unlikely to explain the observed structured representations of color similarity.

### 3.4. Theoretical implications

While the present study found representations of perceptual similarity beyond visual areas, the functional role of these representations still remains unclear. It is possible that higher-order association regions integrate perceptual information with mnemonic information to support subjective experience. In this view, representations in visual areas may primarily reflect the physical structure of sensory input, whereas those in higher-order regions may reflect this information combined with prior experiences and knowledge. The higher-order regions identified in our study, especially the LPFC, overlap with or are functionally connected to the frontoparietal and default mode networks (Figure 4B and Supplementary Figure 8). These networks have been implicated in representing internal information related to prior knowledge (González-García et al., 2018; González-García & He, 2021; Ujhelyi et al., 2026).

Relatedly, subjective similarity judgments have been shown to reflect one’s discrimination capacity, a measure of perceptual resolution in distinguishing stimuli (Moharramipour et al., 2026). This suggests that subjective similarity may involve a readout of information embedded within the visual system, a process that may engage higher-order regions such as the LPFC. Importantly, such a readout may be an automatic component of perception, available even without explicit attention (Lau et al., 2022), consistent with our findings of similarity representations without actively attending to similarity. For example, when we see the color red, we cannot fail to see it as a color that is more like pink than blue, more like orange than silver, and so on, even when we do not explicitly make these pairwise comparisons.

As these ideas remain speculative, further testing would require demonstrating that higher-order association regions more strongly reflect subjective perceptual similarity (e.g., measured by similarity judgment tasks as in this study) than similarity based on physical properties or that inferred from perceptual discrimination capacity (Moharramipour et al., 2026). However, the present study could not effectively address this question, because behaviorally derived subjective similarity showed little inter-individual variability and was virtually indistinguishable from the similarity structure based on the physical properties of the colors. Future studies using a much larger set of colors or more complex stimulus domains, such as faces (Moharramipour et al., 2026), may produce greater inter-individual variability, thereby better isolating subjective perceptual similarity and exploring these hypotheses.

## 4. Materials and Methods

### 4.1. Human fMRI experiment

#### 4.1.1. Participants

Eight participants (19 to 28 years old, mean age of 22; five males, two females, and one who did not disclose their gender) were recruited for this study. Each participant was scanned for a substantial duration. Despite the modest number of participants, this extensive sampling could enable fairly reliable group-level analysis by reducing noise in the estimates from each participant.

Participants were recruited in two cohorts. Some details of the experimental setup differed between the initial cohort (N=4) and the subsequent cohort (N=4). In the following, we primarily describe the setup used for the second cohort, while highlighting key differences for the first cohort. The experimental procedures and sampling/exclusion criteria for the second cohort were pre-registered on the <u>AsPredicted</u> platform. We adhered to our pre-registration when conducting the study for these participants.

The study received approval from the Ethics Review Committee at RIKEN and was conducted in accordance with its ethical guidelines. Informed consent was obtained from participants before the experiment, and they were compensated 6500 yen (approximately 40 US dollars) for each day of participation (roughly 2.5 hours each day).

#### 4.1.2. Experimental design

##### 4.1.2.1. Repetition-suppression (RS) paradigm

This paradigm was designed to allow the extraction of representational dissimilarity between stimulus color pairs within individual voxels by leveraging the RS phenomenon.

The color stimuli were presented in a checkerboard pattern, consisting of a default gray and the intended color, which covered the entire screen (Figure 1A). Checkerboard pieces were 40 × 40 pixels on a 1920 × 1080 screen resolution. There were two types of stimulus presentation blocks: one showing only a single color and the other showing two colors alternating every 250 ms (4 Hz). In both block types, the checkerboard pieces also shifted positions (flickered) every 250 ms according to a predetermined sequence. This sequence was designed to counterbalance the presentation of colors and the default gray across the checkerboard pieces over time. Each stimulus block lasted for 6 seconds, with a random jitter of 1 second. Stimulus blocks were separated by blank fixation blocks of 4, 6, or 8 seconds, with a random jitter of 1 second.

Five colors (see Figure 1B) were used in an experimental run, resulting in 15 conditions: 5 featuring each individual color and 10 featuring all possible combinations of color pairs. Each condition was repeated three times randomly within a run, with each repetition preceded once by the 4, 6, and 8-second blank fixation blocks. We ensured that no condition appeared consecutively. In the color-pair conditions, the choice of the color that appeared first at the start of a block was randomized. The entry point into the flickering sequence was also randomized across all blocks. See Supplementary video for examples of stimuli.

Participants were instructed to fixate on a small white circle (5 pixels in radius on a 1920 × 1080 screen resolution) presented at the center of the screen throughout the run while viewing the stimuli. To enhance their engagement in the task, they were asked to press a key at the beginning of each stimulus block and another key at its end. They were also instructed to press a third key both at the beginning and end of the block when the block included control target colors. The control colors differed significantly in hue and luminance from the experimental colors, making them easily recognizable to participants. A total of four control colors (white, black, bright gray, and yellow) were used, with two randomly selected (with replacement), paired, and utilized in each run. These selected colors were shown to participants immediately before each run, so they knew which control colors would appear in the run. The control color blocks appeared randomly only three times within a run consisting of 48 stimulus blocks in total (i.e., 3 × 15 normal blocks and three control blocks in a run). The control colors were never paired with the experimental colors. At the end of each run, a score was shown to participants, calculated based on their performance (i.e., the accuracy of their key presses and reaction times) to further encourage them to remain alert throughout the run by aiming for a high score. Before each run, participants were instructed to use either their left or right hand to press the keys; all key presses were made with one hand. The use of the left and right hands was counterbalanced across the runs. These key presses were included solely to help participants stay engaged with the stimuli, as simply staring at the stimuli without interaction could become monotonous, leading to drowsiness or even sleep. Before scanning, participants completed a brief practice run outside the scanner to familiarize themselves with the task and instructions. Importantly, color similarity was task-irrelevant, as participants were not required to make use of the similarity between colors in any way.

Four experimental color sets, each containing five colors, were designed and used separately in the above paradigm; all colors were physically isoluminant (Figure 1B), as measured with a spectroradiometer (JETI specbos; JETI Technische Instrumente GmbH). The second cohort completed six runs of each (a total of 4 × 6 = 24 runs), pseudo-randomly distributed across four fMRI sessions. This equated to around 235 minutes of fMRI data recorded during the RS paradigm for each participant. Note that we did not use all colors as a single set but divided them into smaller sets of five colors for practical reasons. A large set contains a substantially greater number of color pairs, making it impractical to sample them all with sufficient repetitions across our sessions.

For the first cohort, two other color sets were used: one containing five non-isoluminant colors and another containing six isoluminant colors (Supplementary Figure 2). The isoluminant set used for this cohort had the same luminance as the colors used for the second cohort and was also relatively similar in color to those colors. Participants in the first cohort were also scanned for a substantial duration while undergoing the RS paradigm across multiple sessions. Specifically, one participant completed 13 runs (eight with set 1, five with set 2; around 160 minutes), another completed 20 runs (ten with set 1, ten with set 2; around 250 minutes), another completed 14 runs (nine with set 1, five with set 2; around 170 minutes), and the remaining participant completed 10 runs (only with set 1; around 115 minutes). Note that the runs with set 2 were longer, because this set contained six colors, resulting in 21 conditions (6 single colors and 15 color pairs). For set 1, 8-second stimulus blocks were used instead of 6-second blocks. Importantly, the overall findings of the study remained similar, and the conclusions did not change, when restricting the analysis to only isoluminant color sets (based on three participants with set 2 from the first cohort and the entire second cohort) (Supplementary Figure 16). However, as this study focused on broad representational similarity, we included the non-isoluminant data in our primary analysis to maximize statistical power and reliability.

##### 4.1.2.2. Similarity judgment task

Participants performed a two-alternative forced-choice (2-AFC) task of judging color similarity. The aim of this task was to quantify the degree of dissimilarity each participant perceives between different pairs of colors.

In each trial, participants were presented with three colors at the center of the screen: one target color at the top, and two other choice colors at the bottom (Supplementary Figure 1). Each color was displayed as a circle with a radius of 145 pixels on a 1920 × 1080 screen. The top target color was presented slightly above the center of the screen, and the two choice colors were displayed below the target, slightly offset to the left and right, such that the three circles formed an isosceles triangular arrangement. Participants were instructed to choose from the two bottom choice colors, the one they perceived as more similar to the top target color. They were required to respond within a 6-second time window by pressing the left or right key with one hand. If they failed to respond within this time window, the trial was skipped and reintroduced randomly later in the run. Upon responding, the chosen color was outlined by a square frame for 200 ms as feedback, followed by a 400 ms blank screen before the next color-trio was presented. All possible color-trio combinations were presented, with the left-right position of the choice colors counterbalanced across repetitions. The same color sets as described in section 4.1.2.1 were used in this task.

As previously mentioned, our primary goal was to estimate perceptual color dissimilarity matrices for each participant based on their 2-AFC responses. In addition to that purpose, we included two control tasks to explore brain regions involved in actively judging color similarity using fMRI. Participants performed 30-second blocks of the 2-AFC color-similarity judgment task, interleaved with two control blocks of the same 30-second duration. The trio of colors was presented as filled circles with slightly varying sizes (a few pixels difference in radius) throughout the run. One control block involved judging similarity based on circle size rather than color, selecting the circle closest in size to the target circle. The other control block did not require any similarity judgment; instead, participants were required to detect a change in the size of a cross fixation presented at the center of the trios (which was also the center of the screen) by pressing either the left or right keys. As in the other blocks, the color-trios changed after every key press. The fixation cross was presented continuously throughout the entire run, across all blocks. Prior to the start of each block, participants were informed of the upcoming block type by a text cue displayed on the screen for 3 seconds. There were no blank intervals between the blocks, except for this text cue.

Participants performed the above task once outside the scanner prior to the fMRI session. The data collected from this preliminary pre-MRI run were used to fine-tune specific task parameters for each participant’s subsequent fMRI run. This included adjusting the degree of variation in circle sizes (which influenced the difficulty of the size-similarity judgment task) and the range within which the fixation-size change occurred randomly relative to trial onset in the no-similarity judgment task. These adjustments were intended to approximately match reaction times and, consequently, the number of trials completed in each control task and in the color-similarity judgment task during the fMRI run.

For the second cohort, the run during fMRI scanning included five blocks of each task type, randomly interleaved such that no block type appeared consecutively. The pre-MRI run included seven blocks for each of the color- and size-similarity judgment tasks and two blocks for the no-similarity judgment task, similarly interleaved so that no block type appeared consecutively. In addition, participants completed another run outside the scanner following the MRI session, consisting of 10 blocks of the color-similarity judgment task only (i.e., a 5-minute run of the color-similarity judgment task only). Note that the task had a fixed duration rather than a fixed number of trials, so participants with faster reaction times completed more trials. Participants completed at least three repetitions (three to four) of all possible color-trios (30 possible trios for a set with five colors) in the pre-MRI run, at least two repetitions (two to three) in the fMRI run, and at least four repetitions (four to eight) in the post-MRI run for the color-similarity judgment task. Therefore, there was ample data (i.e., 2-AFC color-similarity judgments) to reliably estimate the perceptual color dissimilarity matrix for each participant. One dissimilarity matrix was estimated (see section 4.1.3.4) per run (i.e., pre-MRI, fMRI, and post-MRI), and these matrices were averaged to form the final dissimilarity matrix used in the subsequent analysis. There were four color sets (see Figure 1B), each used in a separate session for this task. Within each session, the same color set was used across all three runs of the task (i.e., pre-MRI, fMRI, and post-MRI). Participants were instructed to respond with either their left or right hand inside the MRI scanner, and the hand assignment was counterbalanced across sessions. For the outside scanner runs, participants used their right hand.

For the first cohort, as described in section 4.1.2.1, two different color sets were used (Supplementary Figure 2). Furthermore, instead of the no-similarity judgment control task, these participants completed another control task in which they based their similarity judgments on color luminance (see the non-isoluminant color set in Supplementary Figure 2).

They performed one fMRI run across sessions using only the non-isoluminant color set (with two participants completing two runs). In the pre-MRI run, participants completed 20 blocks of each task type. In the fMRI run, they completed 12 blocks of the color-similarity judgment task and 6 blocks for each of the other control task types. This resulted in participants completing at least eight (eight to thirteen) repetitions per color trio in the pre-MRI run and at least five (five to six) repetitions in the fMRI run for the color-similarity judgment task. As mentioned earlier, the other isoluminant color set was not used during the fMRI scanning for this task in this cohort. Participants performed one run with this set outside the scanner, consisting of 15 blocks of the color-similarity judgment task only (i.e., a 7.5-minute run of the color-similarity judgment task only), allowing participants to complete at least four repetitions of each color-trio. This set had six colors, thus, 60 possible color-trios. Similar to the second cohort, there was ample data to reliably estimate the perceptual color dissimilarity matrix (see section 4.1.3.4) for each color set and participant. One dissimilarity matrix was estimated per run, and then averaged across runs; only the non-isoluminant color set had multiple runs.

Gamma calibration was performed on all screens, and linear RGB values were used to present colors. This procedure ensured consistent color appearance between the runs inside and outside the scanner. However, isoluminance of the colors presented outside the scanner was not verified with a photometer, therefore, unlike the colors presented inside the scanner, isoluminance was not guaranteed. Nevertheless, color appearance is the major determinant of color-similarity judgments, and thus small differences in luminance are unlikely to affect the validity of the assessed perceptual similarity from the runs outside the scanner. Importantly, the dissimilarity matrices derived from the inside- and outside-scanner runs were quite similar and even when we exclusively used the dissimilarity matrices derived from the fMRI runs in our analysis, where available, the findings of the study remained similar.

##### 4.1.2.3. Retinotopy

Participants completed a standard retinotopic paradigm that involved viewing an aperture with an expanding/contracting circular ring and a rotating clock arm, while maintaining fixation at the center of the screen. The goal of this task was to obtain the polarity and eccentricity maps, which were then used to delineate visual areas for each participant. This paradigm was also designed to simultaneously serve as a simple localizer for identifying brain areas responsive to viewing color versus grayscale images, which was relevant to the study. The schematic of the paradigm, along with an example of the estimated polarity and eccentricity maps, is shown in Supplementary Figure 3A. A thorough description of the paradigm and other relevant information is also provided in the legend of this figure. We therefore refer readers to the supplementary materials for more information about the paradigm.

All participants also completed a motion localizer task, which, along with the retinotopic maps, was used to delineate the visual motion area (MT). This task involved viewing stimulus blocks of moving dots interleaved with stimulus blocks of static dots. To maintain participant engagement, participants were required to perform button presses during the task in response to changes in speed, orientation, and density of moving dots, as well as changes in the arrangement of the dots when they were static.

##### 4.1.2.4. Movie watching

Participants watched the first 20 minutes of the *Interstellar* movie during fMRI scanning (the first 15 minutes for the first cohort). Participants were instructed simply to watch the movie. The movie was played without sound and included subtitles. These data were used to perform functional connectivity analysis.

The second cohort completed the movie-watching, retinotopy, and motion localizer tasks entirely during their first scanning session (of five sessions), whereas the remaining four sessions were dedicated to the RS paradigm, with some time allocated to the similarity judgment task. In the first cohort, however, the movie-watching, retinotopy, and motion localizer tasks were distributed across scanning sessions rather than all being performed in one session.

##### 4.1.2.5. MRI apparatus and scanning parameters

MRI data were collected at the Center for Brain Science, RIKEN, using a Siemens Prisma 3T scanner with a 64-channel head coil. Stimuli were displayed on a translucent screen using an MRI-safe projector located approximately 1 m from participants’ eyes, yielding a full horizontal visual angle of approximately 30°. The screen resolution was set to 1920 × 1080, and the MRI room was kept dark during the scans.

All functional images were obtained using a 2D gradient echo-planar imaging (EPI) sequence with the following key parameters: voxel size of 2×2×2 mm³, 60 axial slices, field of view (FOV) of 192 × 192 mm, repetition time (TR) of 1000 ms, echo time (TE) of 30 ms, flip angle of 64°, anterior-posterior (A-P) phase encoding direction, and multiband acceleration factor of 4.

Two additional EPI images with identical spatial geometry (voxel size of 2×2×2 mm³, 60 axial slices, FOV of 192 × 192 mm, TR of 8000 ms, TE of 30 ms, flip angle of 90°) and opposite phase encoding directions (A-P and P-A) were acquired for four TRs each, once at the beginning and once in the middle of each session. These images were later used to correct susceptibility artifacts in fMRI data preprocessing.

Structural images were acquired at 0.7×0.7×0.7 mm³ using a 3D MPRAGE sequence (volume of 290 × 320 × 256 voxels, inversion time (TI) of 1100 ms, TR of 2180 ms, TE of 2.95 ms, and flip angle of 8°) and at 1×1×1 mm³ using a 3D MPRAGE sequence (volume of 256 × 256 × 192 voxels, TI of 900 ms, TR of 1700 ms, TE of 2.73 ms, and flip angle of 8°). The 0.7 mm image was acquired only once for each participant, whereas the 1 mm image was acquired once per session, resulting in multiple 1 mm images per participant. The 1 mm images were combined (using FreeSurfer) and used in subsequent analysis, as their combined image yielded higher quality than the single 0.7 mm image.

#### 4.1.3. Data analysis

##### 4.1.3.1. fMRI data processing

The structural data were processed using the FreeSurfer toolbox’s default pipeline. Susceptibility-induced field distortions were estimated from the acquired A-P and P-A EPI images using the FSL toolbox, and field distortion correction was subsequently performed using the AFNI toolbox. The following preprocessing steps were then applied using the AFNI toolbox: despiking, physiological noise correction (for heartbeat and respiration), slice-timing correction, head-motion correction, registration to the structural image, exclusion of non-brain voxels (brain masking), and normalization of voxel time series to a mean of 100. Head-motion correction and registration were applied in a single interpolation step. Volumes with excessive head motion (frame-to-frame displacement > 0.3 mm) were censored in the subsequent analysis, following AFNI’s default procedure.

The design matrix was constructed using the AFNI toolbox. It included low-order polynomial basis functions and head-motion parameters as nuisance regressors, and consisted of one regressor per experimental condition modelled using AFNI’s dmUBLOCK hemodynamic response function. All runs were concatenated, and the regressors for the same conditions were merged across runs to produce a single regressor per condition per color set, while the nuisance regressors were modeled separately for each run within the design matrix.

Beta estimates in each voxel were obtained using a two-step procedure as follows. First, a second-order autoregressive-moving-average, ARMA(2,2), model was fitted to each voxel’s time series to remove predictable components of the signal, expected to primarily reflect noise and artifacts. This step retained the residual signal not predicted by the general temporal structure, referred to as the innovations. Second, a robust regression was applied to fit the model to the innovations. Robust regression downweights the influence of outlier samples, which may improve the beta estimates. Custom MATLAB code was used to implement this procedure, utilizing MATLAB’s built-in “arima” and “robustfit” functions. Essentially, the ARMA model served as a somewhat aggressive filter that helped denoise the signal (e.g., likely removing complex scanner-related drifts or structured noise); however, at the same time, it could also attenuate stimulus-related responses, alongside reducing noise and artifacts. While this may reduce absolute beta estimates, we thought that it might result in more accurate beta contrasts (i.e., the difference in estimates between conditions), which is the focus of the main analysis (see section 4.1.3.3). We caution that this fitting pipeline is non-standard and is not recommended as a general preprocessing strategy due to the risks described above.

Nevertheless, to ensure that the study’s conclusions were not dependent on the specific model-fitting procedure, we also applied a typical generalized least squares (GLS) similar to that implemented in AFNI for comparison. The final results from both methods were largely consistent (see Supplementary Figure 17 and compare with Figures 3 and 4), although the typical approach yielded slightly lower sensitivity in visual areas such as V2 and V3.

For the second cohort, we pre-registered specific criteria to exclude low-quality fMRI runs from analysis. We also pre-registered that we would exclude a participant entirely from the analysis if they had a notable number of low-quality runs across sessions or entirely missed more than one of their scheduled scanning sessions. However, no participant was excluded, as they successfully completed all their scanning sessions, and the vast majority of runs satisfied our quality criteria. Low-quality runs were defined as those with excessive head-motion or low task attentiveness. No runs across participants were excluded due to excessive head-motion. However, 7% of the RS paradigm runs across participants (7 out of 96 runs; about two runs per participant) were excluded because they met our low-quality criteria based on button-pressing accuracy and reaction times during the run. Five out of sixteen runs from the similarity judgment task across participants (about one to two runs per participant) were excluded for low quality. For this task, quality was evaluated based on response accuracy in the size-similarity judgments and response consistency in the color-similarity judgments during the run. Please refer to our AsPredicted pre-registration for further details. Note that we used a strict data quality criterion, and brief sleepiness and reduced attentiveness during the run were sufficient to make the run meet the exclusion criteria. The first cohort was not pre-registered like above. However, they all performed well attentively during their sessions, and we did not exclude any of their runs from the analysis.

##### 4.1.3.2. Regions of interest

Visual Areas of V1, V2, V3, V4, V3A, V3B, MT, were delineated manually for each participant according to their retinotopic polarity and eccentricity maps (Supplementary Figure 3) derived using the SamSrf Matlab toolbox (https://osf.io/2rgsm/). Delineation of area MT was also guided by a separate motion versus static localizer in combination with the retinotopic maps (see section 4.1.2.3). Another visual ROI in the medial occipital cortex was manually drawn, extending from the outer boundary of the retinotopic eccentricity map to the parieto-occipital sulcus (POS) and anterior calcarine fissure, essentially encompassing the remaining medial occipital cortex not covered by the retinotopically delineated visual areas. We named this ROI the MOC; it included regions commonly associated with the visual periphery as well as regions around the POS, such as V6 and DVT. Higher-order brain areas, including the lateral prefrontal cortex (LPFC), parietal association cortex (PAC), medial frontal cortex (MFC), and orbitofrontal cortex (OFC), were defined by grouping parcels from Glasser’s atlas (Glasser et al., 2016) according to their anatomical locations. When delineating the PAC, we ensured there was no overlap with the manually delineated V3A and V3B. See Supplementary Figure 3B for an example of the above ROIs in a participant.

##### 4.1.3.3. Voxel-level neural RDMs

The neural representational dissimilarity between each pair of stimuli within a voxel (*Di,j*) was estimated as follows:

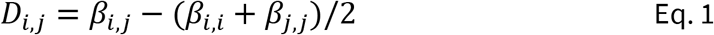

where *β_i,j_* indicates the beta estimate of the voxel during blocks where stimulus *i* and *j* were paired, and *β_i,i_* and *β_j,j_* correspond to the beta estimate during blocks showing only stimulus *i* and only stimulus *j*, respectively. The idea is that if two stimuli have more dissimilar neural representations within a voxel (i.e., less overlap in their neural activity pattern), there will be less RS, and thus a higher fMRI activation level when presented as alternating pairs (Barron et al., 2016). The *β_i,i_* and *β_j,j_*, cases with the most RS, serve as the baseline for evaluating this increased activation level. All pairwise representational dissimilarity (*Di,j*) was calculated as above, and the RDM was derived for each individual voxel and for each of the color sets. See Figure 2A,1 for the RDM of an example voxel in color set 1.

The neural RDM may contain cells with negative values (see Figure 2A,1 example), as the equation above allows. If neurons within a voxel encode colors, the voxel’s RDM is expected to have positive cells. However, due to fMRI noise, some cells can still take on negative values. If neurons in a voxel show no response (or an invariant response) to color, the resulting RDM is expected to contain both negative and positive values in a noisy manner. Thus, the positivity of the RDM, evaluated as follows, could indicate whether a voxel is responsive to color.

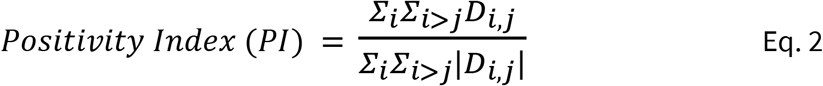

This positivity index is simply computed by taking the sum of the RDM cells and dividing it by the sum of their absolute values. This index ranges from -1 to 1, with higher values indicating the presence of color-related responses. The positivity indices computed from the RDMs corresponding to different color sets were averaged to form the final positivity index value for the voxel.

##### 4.1.3.4. Perceptual behavioral RDMs

The perceptual dissimilarity matrix, referred to as the behavioral RDM, was obtained for each color set from 2-AFC judgments participants made in the color-similarity judgment task (see section 4.1.2.2 and Supplementary Figure 1). Each cell in the behavioral RDM represents the perceived dissimilarity between a pair of colors.

A machine-learning approach was used to estimate the behavioral RDM from the 2-AFC judgments as follows. Two-dimensional embeddings were first randomly initialized. In each iteration, the embeddings were updated using a loss function that penalized mismatches between the similarity judgments predicted from the embeddings (based on the Euclidean distances between embedded color points) and the participant’s actual similarity judgments (Hojjati et al., 2025; Moharramipour et al., 2026). An elastic regularization was incorporated into the loss function to control the scale of the embeddings and prevent overfitting. Once the embeddings were obtained through this machine-learning approach, the RDM was derived by calculating the Euclidean distances between the embedded color points. Finally, the RDM was normalized by dividing it by its maximum value. One RDM was estimated from each task run (see section 4.1.2.2) and then averaged across runs to form a single RDM; thus, one behavioral RDM was obtained for each color set and participant (see Figure 2B,1 for an example behavioral RDM).

##### 4.1.3.5. Repetition-suppression multi-voxel-pattern-analysis (RS-MVPA)

We performed our main analysis at the ROI level by incorporating an MVPA approach. First, we aimed to identify ROIs that contained voxels exhibiting general color-related responses. Second, and more importantly, we sought to identify ROIs containing voxels that represent color similarity consistent with perceptual similarity. In both cases, we used a split-half pattern correlation as a form of MVPA (Haxby et al., 2001) as follows.

For the first aim, we divided the data into two halves: one containing the odd runs and the other containing the even runs. We then ran our model-fitting separately on each half, resulting in one neural RDM per voxel and per color set for each half. The positivity index (see section 4.1.3.3) was calculated for each voxel and color set, then averaged across color sets, yielding one positivity index value per voxel for each half of the data. Next, we assessed the consistency of the positivity index values of voxels within each ROI between the even and odd observations (Figure 2A, 3). This consistency was quantified by arranging voxel values into a vector and computing the Pearson correlation between the even and odd vectors. The correlation coefficient was then transformed into a z-value using Fisher’s z-transformation for a subsequent group-level t-test (one-tailed t-test, as negative correlations are theoretically meaningless). In summary, each ROI was assigned a z-value for each participant, indicating the extent to which it exhibited a pattern of voxels with consistent color-related responses.

For the second aim, similar to the above, one neural RDM was first computed per voxel and per color set, separately, for even and odd data. To assess how similar a voxel’s RDM is to the actual perceptual RDM, cosine similarity was employed. The upper-triangular part of the neural and behavioral RDMs was vectorized, and cosine similarity was computed between them (Figure 2B,1). The cosine similarity values obtained for each color set were then averaged to reflect the overall alignment between neural and behavioral RDMs at each voxel. Next, Pearson pattern correlation was computed within each ROI between the cosine similarity values of voxels from the even and odd observations (Figure 2B,2). To evaluate whether this pattern correlation is meaningful—that is, whether the ROI truly contains voxels with representations of perceptual color similarity—a nonparametric permutation test was performed as follows: The above procedure was repeated 10,000 times using randomly permuted behavioral RDMs where rows and columns were simultaneously shuffled (Mantel, 1967). The pattern correlation values acquired across 10,000 permutations served as the null distribution for evaluating the significance of the actual pattern correlation value (Figure 2B,3). Basically, the idea is that if an ROI truly contains voxels with representations of perceptual color similarity, the pattern correlation is expected to be highest when using the actual behavioral RDMs compared to when using permuted behavioral RDMs. This is because the cosine similarity values of the relevant voxels are expected to be higher and more consistent across observations when computed using the true behavioral RDMs, thereby increasing the pattern correlation. For each ROI and each participant, a z-value was calculated by comparing the actual pattern correlation against its null distribution, specifically, by subtracting the null distribution’s mean from the actual value, and then dividing by the null distribution’s standard deviation. Finally, these participant-level z-values were used in a group-level t-test (one-tailed, as negative values are theoretically meaningless in this context) to assess statistical significance for each ROI.

We validated the above approach through simulation (see Supplementary Figure 5 for details). The simulation demonstrated that our method can identify ROIs containing voxels with representations of perceptual similarity while effectively controlling false positives (i.e., type I error; ROIs without such representations). It remained reasonably sensitive even where only a small subset of voxels within an ROI contained representations of perceptual similarity. Additionally, the simulation showed that the method performs substantially better with cosine similarity than with Pearson-based similarity (i.e., Pearson correlation), a commonly used alternative for measuring similarity between RDMs. With cosine similarity, the method remained significantly more sensitive at lower SNRs (e.g., those expected in experimental data) while effectively controlling type I error, than with Pearson-based similarity. The use of cosine similarity was also theoretically motivated from the outset because values near 0 in the RDMs are interpretable, corresponding to minimal representational dissimilarity. Moreover, cosine similarity incorporates both the sign information (e.g., positivity of the RDMs) and the underlying geometric structure, which is desirable in our case. Cosine similarity is also a widely used metric for comparing two RDMs (Diedrichsen et al., 2020; van den Bosch et al., 2025).

To be critical, our approach for identifying ROIs with a general color response (Figure 2A), unlike our approach for identifying ROIs with representations of perceptual similarity (Figure 2B), may be susceptible to false positives because it lacks a null distribution derived directly from empirical data. For example, factors unrelated to color response, such as the anatomical positioning of voxels within sulci and gyri (i.e., more signal in gyri than in sulci), could artificially inflate the pattern correlation value. In addition, spatial smoothness may inflate the derived z-values. However, these concerns are likely mitigated by the subsequent group-level t-test, which assesses consistency of the effect across participants. Even so, we cannot dismiss the possibility that the positivity index could potentially become high if a voxel responds more strongly to the presence of two colors, regardless of color identity (e.g., reflecting stimulus counting), or become low if it responds less strongly to a stimulus pair in general. This factor could also inflate the pattern correlations and lead to false positives. Nevertheless, the aim of this approach (Figure 2A) was to identify ROIs potentially exhibiting color-related responses, and then to have the second approach (Figure 2B) identify the ROIs that contain representations of perceptual similarity.

All the analysis above was conducted in native surface space. When computing the pattern correlation, a weighted Pearson correlation (Bailey et al., 2018) was used, with the vertex surface areas serving as weights. When projecting data from volume to native surface space (using the FreeSurfer toolbox), given our fMRI resolution, nearby vertices were often assigned the same value corresponding to the same voxel (e.g., 1600 vertices in the right V4 of a participant, with 300 of them being unique). Accordingly, only the vertices with unique values and their corresponding total surface area were included in the computation of the weighted Pearson correlation. Although all main analyses were performed in surface space, we used the term ‘voxel’ for simplicity throughout the manuscript, as the distinction from ‘vertex’ is not critical for conveying the findings.

Lastly, we note that our RS-MVPA analysis can offer higher sensitivity than a comparable univariate analysis, as it is performed at the ROI level. However, this advantage is particularly evident in higher-order brain areas, where relevant voxels may form smaller and sparser clusters, with greater inter-individual variability in their exact spatial locations compared to areas like the visual cortex (see Figure 2A,3 for an example).

### 4.2. Non-human primate electrophysiology experiment

#### 4.2.1. NHP subject

One male rhesus macaque monkey (*Macaca mulatta*) participated in this study (10 years old, 9.5 kg). No food restrictions were imposed throughout the study. The monkey was fed twice daily, with the water supply restricted only on recording days. The experiment was conducted at the Nonhuman Primate Research Center at East China Normal University. All aspects of the study, including animal care, experimental protocols, and surgical procedures, fall within a protocol previously approved by the Institutional Animal Care and Use Committee (permission code: M020150902-2018) at East China Normal University.

#### 4.2.2. NHP experimental apparatus, eye tracking, and task

The monkey sat in a custom-manufactured plexiglass chair with its head restrained inside a lead-shielded recording booth. The monkey’s eyes were positioned approximately 58 cm from a 31.5-inch monitor screen (LG Electronics, Model 32GP83B-B), corresponding to a horizontal visual angle of about 62°. The monitor was set to a resolution of 2560 × 1440, and the booth was kept dark. A tube was positioned beneath the monkey’s mouth and connected to an automated system for reward delivery.

Gaze position was sampled at 1000 Hz from the right eye using an EyeLink 1000 Plus infrared eye-tracking system (SR Research Ltd.). The illuminator module and the camera were positioned above the monkey’s head. An angled infrared mirror was used to capture and re-coordinate monkeys’ eye positions. Standard calibration and validation procedures were performed before each experimental run.

The same RS paradigm used in the human study (Figure 1A) was employed, with a few modifications. The stimulus blocks were 4.5 s long with a random jitter of 0.5 s, and the blank gray blocks were 4 s long with a random jitter of 0.5 s. The monkey was required to maintain central fixation throughout the stimulus presentation and received a liquid reward (∼1.5 ml of orange juice) for successfully holding fixation, which was delivered over a 0.3–1.3 s window during the subsequent blank block. If the monkey made a saccade during the stimulus block, the trial was immediately terminated, and followed by the blank block with no reward delivered. Unlike the human study, the central fixation point only appeared during the stimulus blocks and was absent during the blank blocks. Therefore, we allowed a maximum of 0.5 s for the monkey to move its gaze to the fixation point at the start of stimulus blocks. If the gaze was not on the fixation point by the end of this time, the trial was immediately terminated. In fact, the monkey typically took about 0.2 s to bring its gaze to the fixation point after the stimulus onset. The fixation point was made three times larger (20 pixels radius on a 2560 × 1440 screen resolution) compared to the human version of the task (5 pixels radius on a 1920 × 1080 screen resolution) to make it easier for the monkey to hold fixation. The checkerboard pieces were scaled proportionally to the screen resolution so that they occupied the same relative size on the screen across setups.

Only the color sets 1 and 2 (Figure 1B) were used in this NHP version of the study. As in the human version, each color set was used separately in a run. Each condition (10 color-pair blocks and 5 single-color blocks) was presented randomly in a run while ensuring that no condition appeared twice in a row. The run continued until six successful repetitions per condition were collected, in which fixation was successfully maintained throughout the stimulus presentation.

We ensured that all colors were physically isoluminant using a photometer (SM208; Shenzhen Sanpo Instrument Co., Ltd.). Although we used the same color sets across both studies, the linear RGB values were slightly adjusted in the NHP setup to maintain isoluminance because the screen gamma characteristics were not identical across setups. The absolute luminance of the colors in the NHP setup was higher (65 *cd*/*m*^2^) than in the fMRI human setup (48.5 *cd*/*m*^2^) because the screen used in the NHP setup had a higher achievable contrast. However, the perceived lightness in both setups was consistent (L* = 55.7 in the NHP and L* = 54 in the human fMRI setups). Overall, despite these differences, the color appearances remained visually near-identical across setups.

#### 4.2.3. NHP recording site

A 32-channel microdrive and chamber system (Gray Matter Research) was implanted over the left medial posterior parietal cortex of the monkey (center of the chamber was at anterior-posterior (AP) = -16.9 mm, medial-lateral (ML) = -5.1 mm, 7° to the left and 48° to the posterior). Surgical procedures were described in detail elsewhere (Wang et al., 2024).

The inter-electrode spacing was 1.5 mm. Penetration depth varied slightly across electrodes, with a typical depth of about 8.5 mm (SD = 0.75 mm) from the chamber base. Each electrode penetrated approximately 15 μm deeper per session over nine recording sessions (a total advancement of approximately 0.135 mm from the first to the last session).

A structural T1 image was acquired post-surgery at 0.9×0.9×0.9 mm³ resolution on a 3T Siemens Prisma scanner using a 3D MPRAGE sequence (volume of 256 × 256 × 192 voxels, TI of 1100 ms, TR of 2300 ms, TE of 2.33 ms, and flip angle of 8°). The T1 image was resliced using 3D Slicer software along the chamber’s orientation to facilitate the localization of electrodes. Example T1 slices with the recording site highlighted are shown in Figure 6A. Electrodes were then marked on the T1 image based on their grid locations and penetration depths using 3D Slicer software. Next, the D99.v2.0 macaque atlas (Saleem et al., 2021) was coregistered on the T1 image to identify the anatomical landmarks corresponding to each electrode. Electrode-landmark correspondence was further confirmed by manual inspection using major sulci as anatomical anchors.

The majority of electrodes were located anterior to the POS, with most in areas V6A and MIP, and a smaller subset in area PEc. Several electrodes were also positioned posterior to the POS in area V2 (see Figure 6). Some electrodes were located near or between two anatomical regions and are therefore, in Figure 6, depicted as crossing both regions to reflect uncertainty in their exact anatomical location.

#### 4.2.4. Electrophysiological data acquisition and preprocessing

Nine experimental daily sessions were conducted. In each session, the task was run separately for color sets 1 and 2, for a total of 15 runs per color set across all sessions. Electrophysiological signals from the microdrive were amplified by an Intan RHD2000 and streamed to a SmartBox acquisition system (NeuroNexus) for recording at a sampling rate of 30 kHz. Signals were referenced to stainless steel screws implanted through the skull and in contact with the dura.

LFPs were derived from the electrophysiological data using the following preprocessing steps. Data were notch-filtered, epoched relative to stimulus onset, and downsampled to 500 Hz. A high-pass filter with a 1 Hz cut-off was then applied to remove slow drifts and offsets. Lastly, epochs containing high-amplitude transients (likely movement-related or electrical artifacts; only a few epochs across all runs) and epochs in which gaze fixation was not successfully maintained (see section 4.2.2) were excluded. All these preprocessing steps were performed using the FieldTrip toolbox in MATLAB.

#### 4.2.5. LFP data analysis

Time-frequency (TF) representations were computed from the LFP signal for each trial epoch using a sliding-window Fourier transform with a Hanning taper, implemented by the FieldTrip toolbox in MATLAB. The sliding windows were frequency-dependent and contained 6 cycles at each frequency. Power estimates were derived every 10 ms for frequencies up to 150 Hz in 1 Hz steps. For each trial, the TF map was z-normalized across time relative to a pre-stimulus baseline period (-1.5 s to -0.1 s), separately for each frequency. This pre-stimulus period corresponded to a time period after reward delivery and before stimulus onset.

Figure 5A shows the baseline-corrected (z-normalized) TF, averaged across all trials (about 1,350 trials) in color set 1, for one example electrode in area V2. As shown in this Figure, stimulus-evoked power was predominantly observed in the gamma band (30-80 Hz). The spectral centroid (i.e., the weighted average frequency at each time point, including only z-values above zero) within the gamma-band spectrum was used as a proxy for the dynamics of the local neural population. Specifically, shifts toward higher spectral centroids could reflect elevated excitatory-to-inhibitory drive (Jia & Kohn, 2011; Magazzini & Singh, 2018; Ray & Maunsell, 2010). The spectral centroid pattern was consistent with the stimulus structure and the expected RS effect (Figure 5B), further supporting the use of spectral centroid as a measure of local neural population engagement.

The spectral centroid decreased and reached a relatively steady state at about 1.5 s after stimulus onset (Figure 5B). We therefore used the median spectral centroid value during the steady period to reflect the summary response. This value was computed separately for each of the color-pair conditions and each of the single-color conditions (see the example in Figure 5C). Specifically, for each condition, the trial-wise baseline-corrected TFs (about 6 × 15 = 90 trials) were averaged, and the spectral centroid and its median value were computed over the 1.5–3.5 s period following stimulus-onset. These values were then used in Eq. 1, as in our fMRI analysis, to compute the RDM (Figure 5C). One RDM was derived per color set for each electrode. These RDMs were subsequently compared with reference RDMs based on the RGB color profiles (i.e., the Euclidean distance between gamma-encoded RGB values). For each electrode, cosine similarity between the neural RDM and the reference RDM was computed (similar to Figure 2B,1) separately for the two color sets and then averaged across them. This procedure was repeated (10,000 times) using permuted reference RDMs to find the null distribution of this averaged cosine similarity value (Mantel’s test). Subsequently, the z-value and the *p*-value (one-tailed, as the negative tail is not theoretically expected) were derived for each electrode, and FDR correction was applied across electrodes (Figure 6C). We note that the outcomes remained similar when the L*a*b* color profiles were used to make the reference RDM. We, however, preferred the RGB space as it reflects the stimulus-defined color values and thus could serve as a somewhat perceptually neutral RDM, whereas the L*a*b* space is adjusted to human perception. Lastly, as in our fMRI analysis, we also computed the positivity index (Eq. 2; see section 4.1.3.3) and averaged the value across color sets for each electrode (see Figure 6B). If an electrode contains color-related information, this value is expected to be high, approaching 1.

## Acknowledgements

We thank Nikolaus Kriegeskorte, Bevil Conway, Tomoya Nakamura, Aurelio Cortese, Qi Lin, Joshua Tan for their helpful comments and suggestions on the manuscript.

## Data and code availability

The raw data, processed data, and the analysis code are publicly available in the following repository: https://neurodata.riken.jp/r/Shibata/Perceptual%20similarity%20representations%20beyond%20the%20sensory%20cortex/ 

## Author Contributions

**Ali Moharramipour**: Conceptualization, Methodology, Software, Investigation (human data collection), Formal analysis, Visualization, Data curation, Project administration, Writing– original draft, Writing–review and editing

**Zhiyong Jin**: Investigation (monkey data collection), Writing–review and editing

**Ning Su**: Investigation (monkey data collection)

**Yijun Ge**: Software, Writing–review and editing

**Takeru Matsuda**: Methodology, Writing–review and editing

**Kazuhisa Shibata**: Resources, Writing–review and editing

**Sze Chai Kwok**: Funding acquisition, Resources, Project administration, Writing–review and editing

**Hakwan Lau**: Conceptualization, Methodology, Funding acquisition, Resources, Project administration, Supervision, Writing–review and editing

