## Supplementary materials for "Perceptual similarity representations beyond the sensory cortex"

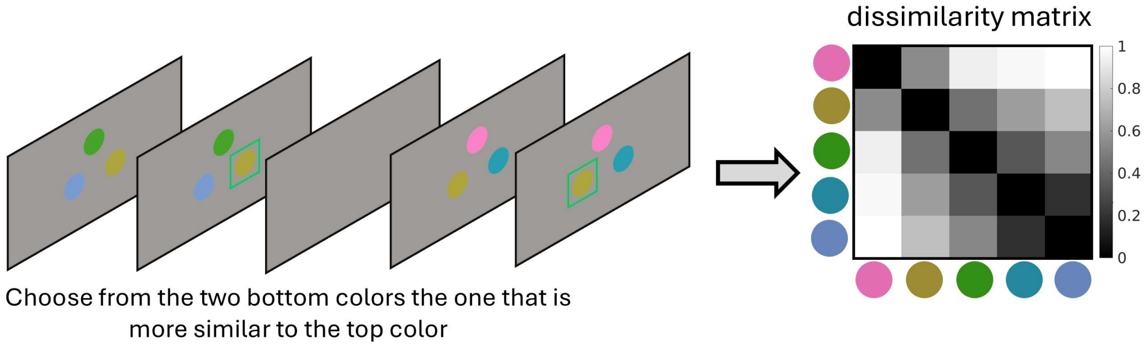


**Supplementary Figure 1. Behavioral color-similarity judgment task.**
In each trial, participants were presented with three colors: one at the top as the target color and two other colors at the bottom. They were required to choose the bottom color that they perceived as more similar to the top target color. They expressed their response by pressing the left or right key. Upon responding, the chosen color was outlined by a green square frame for 200 ms as feedback, followed by a 400 ms blank screen before the next color trio was presented. All possible trio color combinations were presented multiple times, with the left–right position of the bottom colors counterbalanced across the repetitions. This illustration is for explanatory purposes and does not reflect the actual proportions presented on the screen during the task.

For each participant, the dissimilarity matrix, which reflects their perceived dissimilarity between colors, was constructed from their responses across trials using a machine-learning approach (see section 4.1.3.4). Participants performed the color-similarity judgment task both inside and outside the MRI scanner, and the resulting dissimilarity matrices were averaged and used in the main analysis as the behavioral RDMs.

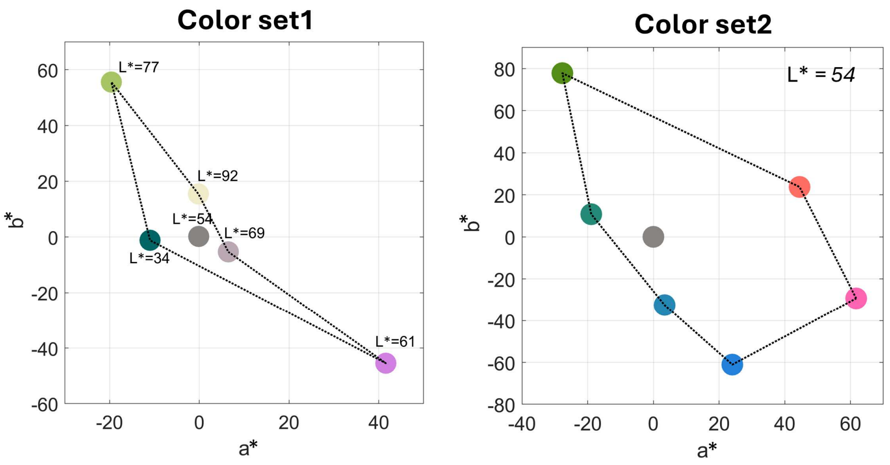


**Supplementary Figure 2. The color sets used in the experiment for the first cohort.**
The first color set contained five non-isoluminant colors, and the second color set contained six isoluminant colors. The L*a*b* values shown were measured by a spectroradiometer.

We note that the overall findings of the study remained similar and the conclusions did not change when restricting the analysis to only isoluminant color sets (based on three participants with set 2 from the first cohort and the entire second cohort; see section 4.1.2.1) as shown in Supplementary Figure 16. However, as this study focused on broad representational similarity, we included the non-isoluminant data in our primary analysis to maximize the power and the reliability of the study.

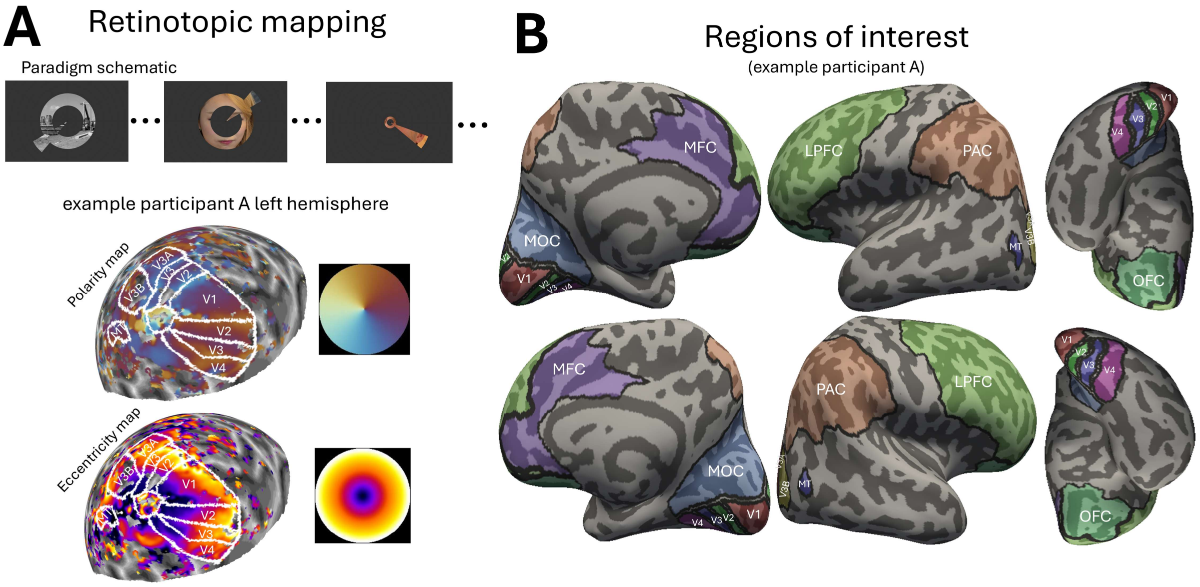


**Supplementary Figure 3. The retinotopic paradigm and the regions of interest.**

**(A)** Participants viewed an aperture containing a circular ring that gradually expanded or contracted, and a clock arm that gradually moved clockwise or counterclockwise every second (i.e., at each scanning TR). The circular ring covered eccentricity levels from 0° (center) to 8.5°, and the clock arm spanned the full polar angle (0° to 360°) in 10° steps multiple times (24 cycles for eccentricity and 12 cycles for polarity in a run). After every 108 seconds, which included three full polar angle cycles (either clockwise or counterclockwise) and simultaneously six eccentricity cycles (either expanding or contracting), a 25-second blank fixation period was presented. This sequence was repeated four times to cover all 2 × 2 combinations of expanding and contracting eccentricity, and clockwise and counterclockwise polar-angle directions. Three image categories, consisting of faces, non-faces (e.g., scenes, objects), and scrambled images, were displayed within the aperture, either in color or grayscale, in 6-second blocks, with the exemplar image changing every 500 ms. The presentation of image categories in color or grayscale (3-by-2 factorial design) was counterbalanced with respect to each other (i.e., an equal number of colored and grayscale faces, non-faces, and scrambled image blocks) and with respect to the aperture's circular ring and clock arm status. Therefore, with this paradigm, we could obtain not only the eccentricity and polarity maps but also other localizers, especially the color-versus-grayscale localizer relevant to this study. Participants were instructed to fixate on the black dot at the center of the screen the entire time. They were also told to press a key whenever the black dot changed color to red, which helped prevent them from shifting their gaze away from the center. Every 200 ms, there was a 2.5% chance that the black dot would change to a random color, including red. To further engage them, they were instructed to press another key every time the clock arm reached 0°, 90°, 180°, or 270°. The second cohort completed three runs of this retinotopic paradigm, totaling 27 minutes. For the first cohort, no specific image category (i.e., random images from scenes, objects, or faces) was presented in either color or grayscale, alternating every 20 seconds (i.e., 20-second blocks of colored or grayscale images), with the exemplar image changing every 500 ms. These participants completed three runs of the task, except for two who completed four runs.

**(B)** Regions of interest in one example participant are visualized. For each participant, V1, V2, V3, V4, V3A, and V3B were delineated based on their polarity and eccentricity maps (see panel A for the corresponding maps of the example participant shown in this panel). Area MT was delineated using these maps, along with a separate motion localizer map (participants completed two runs of a motion localizer, which included stimulus blocks of motion dots and static dots). MOC was manually drawn to encompass the medial occipital regions not covered by the retinotopically delineated ROIs. PAC, LPFC, MFC, and OFC were defined by grouping parcels from Glasser's atlas (Glasser et al., 2016) based on their anatomical locations.

LPFC included the following Glasser's atlas parcels: FEF, PEF, 55b, 8Av, 8Ad, 9m, 8BL, 9p, 10d, 8C, 44, 45, 47l, a47r, 6r, IFJa, IFJp, IFSp, IFSa, p9-46v, 46, a9-46v, 9-46d, 9a, 10v, a10p, 10pp, 6a, i6-8, s6-8, p10p, p47r.

MFC included the following Glasser's atlas parcels: 33pr, p24pr, a24pr, p32pr, a24, d32, p32, 8BM, 10r, 25, s32, a32pr, p24.

OFC included the following Glasser's atlas parcels: 47m, 11l, 13l, OFC, 47s, pOFC.

PAC included the following Glasser's atlas parcels: PGp, PGs, PGi, PFm, PF, PFt, PFop, IP0, IP1, IP2, LIPv, LIPd, VIP, AIP, MIP, 7PC, 7AL, 7Am, 7PL, 7Pm.

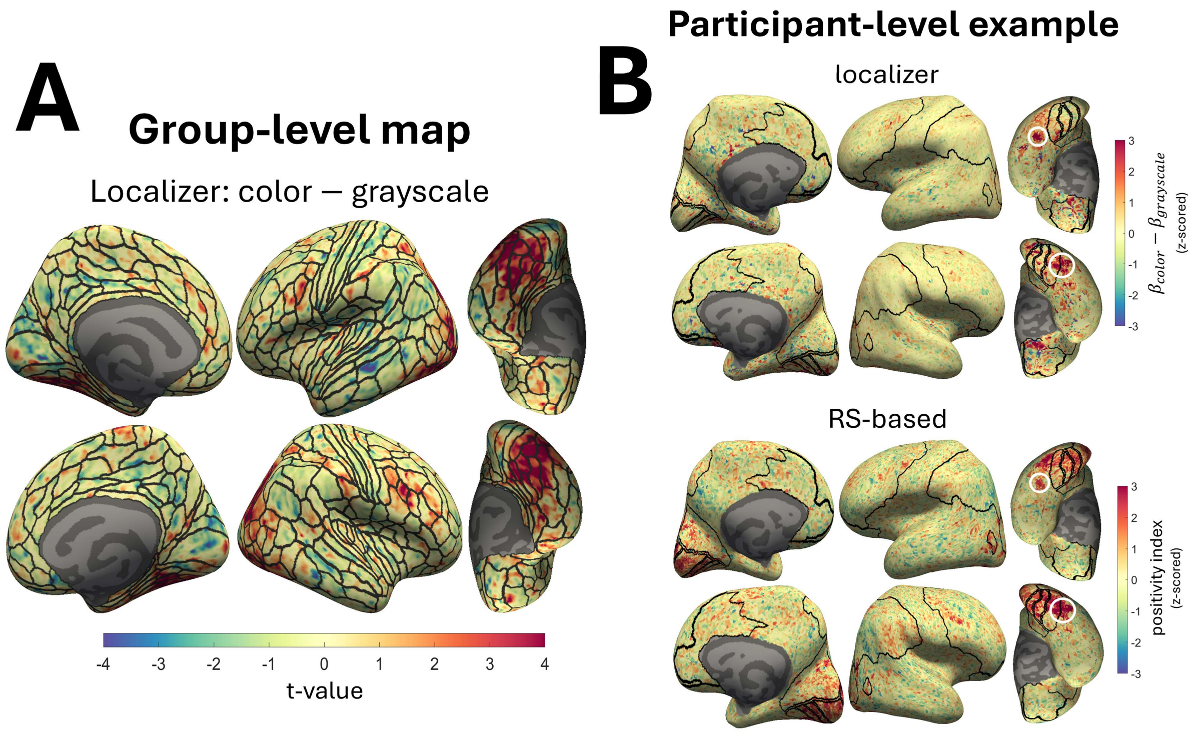


**Supplementary Figure 4. An exploratory color localizer.**

**(A)** This panel shows the group-level (N = 8) contrast map of viewing colored versus grayscale images displayed in the retinotopic paradigm (Supplementary Figure 3A). The color bar represents the t-value from the group-level t-test applied to the beta contrast values ($\beta$-color – $\beta$-grayscale; design matrix included one regressor per block type). The individual participants' beta estimates were projected onto an average brain (Freesurfer's fsaverage), with 5 iterations of surface smoothing applied both before and after projection as implemented in FreeSurfer's mri_surf2surf tool, prior to performing the group-level test. AFNI's afni_proc.py tool was used to perform model fitting and obtain beta estimates for each individual, incorporating the same preprocessing steps and nuisance regressors described in section 4.1.3.1. **(B)** The color-versus-grayscale ($\beta$-color – $\beta$-grayscale) from the retinotopic paradigm and the positivity index from the RS paradigm (Figure 2A,2) are visualized in the top and bottom panels, respectively, for an example participant (no spatial smoothing was applied when projecting the maps onto the native surface). For better comparison, the values in both maps are z-scored using the mean and standard deviation across all vertices. All maps, including panel A, are displayed unthresholded and are intended for descriptive visualization. The ROIs outlined in black are the same as those shown in Supplementary Figure 3B. The white circles indicate areas with particularly strong effects in the color-versus-grayscale localizer. These regions also consistently showed strong RS effects (i.e., high positivity index). However, there are regions with notable effects in the RS-based map without apparent effects in the localizer map. This may indicate greater sensitivity of the RS-based approach for identifying color-responsive areas. That said, this comparison may not be entirely fair because the RS-based approach included substantially more data.

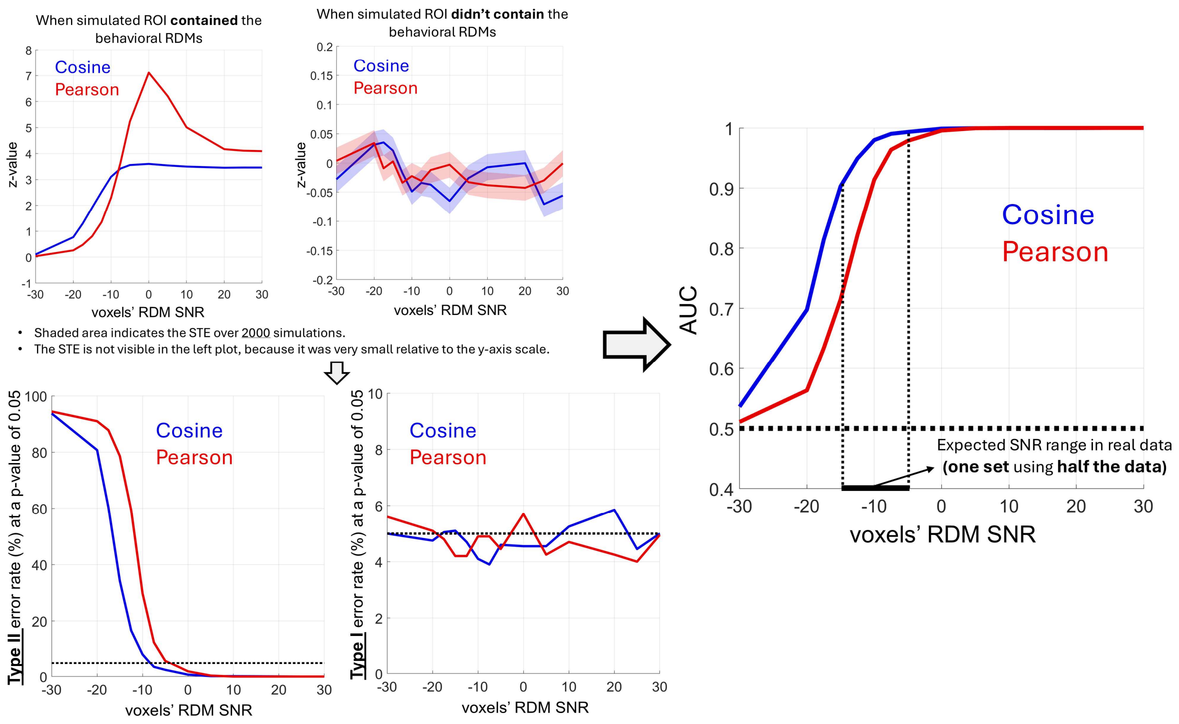


**Supplementary Figure 5. Simulations to validate the RS-MVPA approach (see Figure 2B).**
An ROI was simulated with voxels containing perceptual behavioral RDMs (as in the main study, four 5×5 behavioral RDMs were assumed; see Figure 1B). Gaussian noise with zero mean was added to voxel RDMs to simulate even and odd observations. Various signal-to-noise ratios (SNRs), expressed in decibels (dB), were examined, with 2000 simulations conducted at each SNR level. The proportion of voxels in the ROI with the behavioral RDMs was randomly varied between 1% and 60% across simulations. The remaining voxels without the behavioral RDMs were assumed to have purely noisy RDMs (zero-mean Gaussian noise), indicating noisy activations. Notably, simulation outcomes remained similar even when these remaining voxels were assigned random RDMs that were consistent between even and odd observations rather than purely noisy RDMs. In addition to the cosine similarity used in the study, Pearson-based similarity (i.e., Pearson correlation) was also evaluated for comparison. As with the real data, voxel RDMs are allowed to have negative cells (with diagonal cells always zero). However, when a voxel contains behavioral RDM, all cells are positive, though added noise may still cause cells to become negative.

The top-left plots show the mean z-value across simulations where the ROI contained voxels encoding the behavioral RDMs (left panel) and across null simulations where the ROI did not contain the true behavioral RDMs but a different (permuted) behavioral RDMs (right panel). The results (right panel and Type I error) remained similar when assuming noise RDMs for all voxels in the null simulation rather than different (permuted) behavioral RDMs. The bottom-left plots illustrate the Type II and Type I errors at a *p*-value of 0.05. The right plot shows the Area Under the Curve (AUC) of the Receiver Operating Characteristic (ROC) curve, indicating the overall performance of the RS-MVPA approach at various SNR levels. The approach using both cosine and Pearson similarities effectively controlled false positives (i.e., Type I error), but it was noticeably more sensitive when cosine similarity was used. Given the expected SNR range of our fMRI data (-15 dB to -5 dB for each color set and each even and odd data), the use of cosine similarity appears crucial due to its superior performance at lower SNRs.

While the proportion of voxels with behavioral RDMs in an ROI was randomly set between 1% and 60% across simulations (e.g., hypothetically as low as 1% in non-visual areas, and as high as 60% in a visual area, such as V4), the approach remained reasonably sensitive even when simulations were restricted to ROIs with a low proportion of voxels with behavioral RDMs. For instance, with a proportion of 1% to 5%, the AUC ranged from 0.60 to 0.93, and with a proportion of 5% to 10%, it ranged from 0.74 to 0.99, across an SNR range of -15 dB to -5 dB using cosine similarity. These AUCs were substantially lower when Pearson-based similarity was used. We note that the ROI was assumed to contain 500 voxels across simulations, roughly matching the size of visual areas such as V1 to V4.

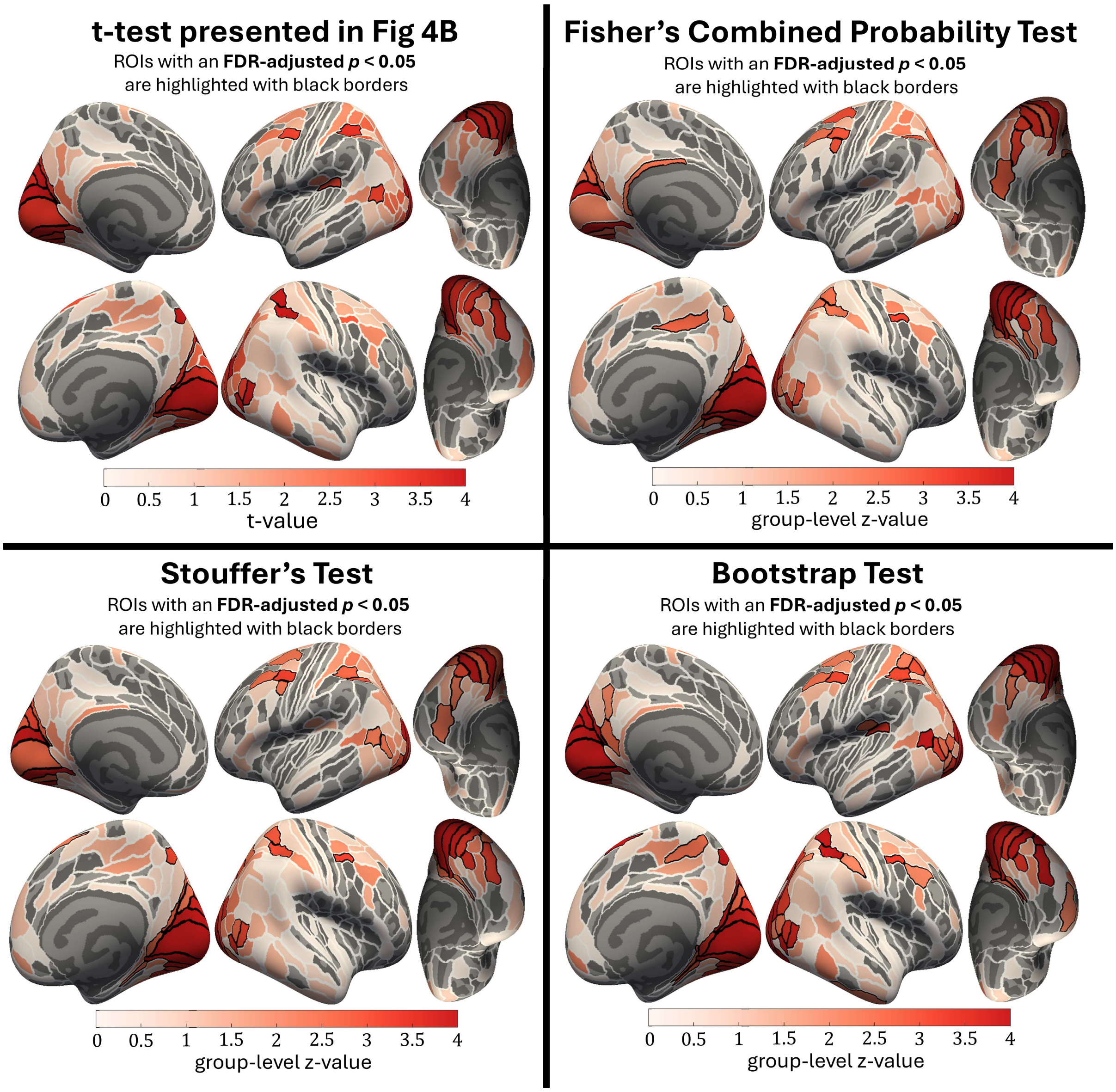


**Supplementary Figure 6. Same as Figure 4B, but using alternative approaches to obtain group-level statistics.**

The top-left panel is identical to Figure 4B and shows the results from a t-test on individual-level z-values. The top-right and bottom-left panels display the results using Fisher's combined probability test and Stouffer's test, respectively, both of which combine the evidence from individual-level *p*-values/z-values to derive the group-level significance. The bottom-right panel shows the results from a bootstrap test that assesses whether the group-mean z-value is significantly above zero.

Collectively across these tests, several areas surrounding the intraparietal sulcus (AIP, 7PC, MIP, LIPv, IPS1, IP0, IP1, IP2, VIP, PFt) and the posterior LPFC (55b, PEF, 8Av, 8C, 6a) in either hemisphere survived the FDR correction.

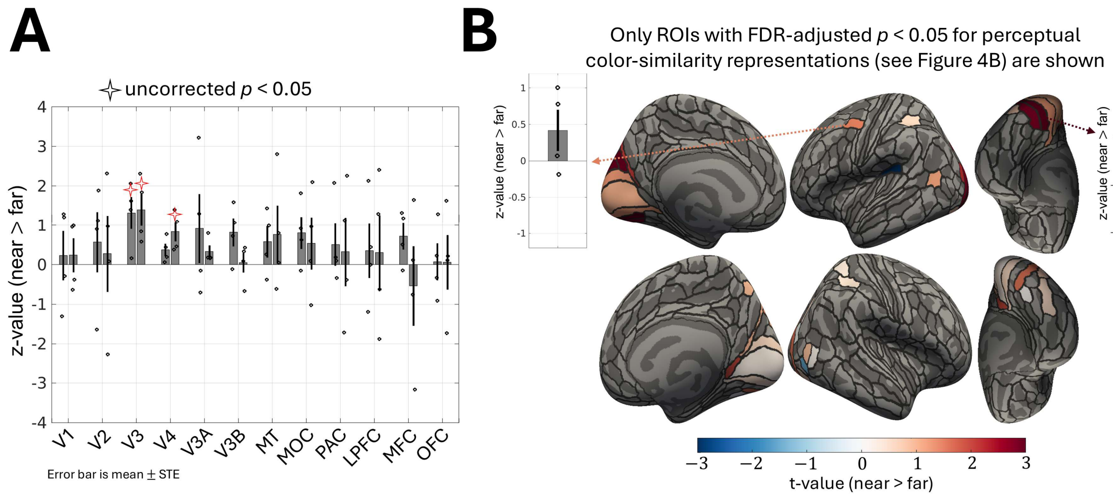


**Supplementary Figure 7. Fine-grained versus coarse-grained representations.**

**(A)** For the second cohort (N = 4), two color sets were systematically used to contain near colors (sets 2 and 4; Figure 1B) and two other color sets to contain far colors (sets 1 and 3; Figure 1B). Voxels whose representations are aligned with near color sets would likely require finer-grained neuronal color coding, whereas for far color sets, coarser-grained coding could be sufficient. A similar RS-MVPA analysis to Figure 2B was applied, but focused on the contrast between the ROI-level measure derived from the near and far sets. Most areas, including the PAC and LPFC, exhibited a trend toward containing voxels that better represent near than far color sets, with this trend being particularly pronounced in V3 and V4 (i.e., only V3 and V4 showed a two-tailed uncorrected $p < 0.05$ from a t-test applied to individual-level z-values). **(B)** Same analysis as panel A, but focusing on Glasser's atlas ROIs that survived FDR correction in the RS-MVPA analysis (see Figure 4B). The color map shows the t-values of a t-test applied to individual-level z-values. Some visual areas (e.g., V3, V4) and the ROI in the left LPFC showed a trend toward better representations of perceptual similarity for near colors.

Notably, because the RDM geometries were not exactly matched between the near and far sets, these results should be interpreted with caution.

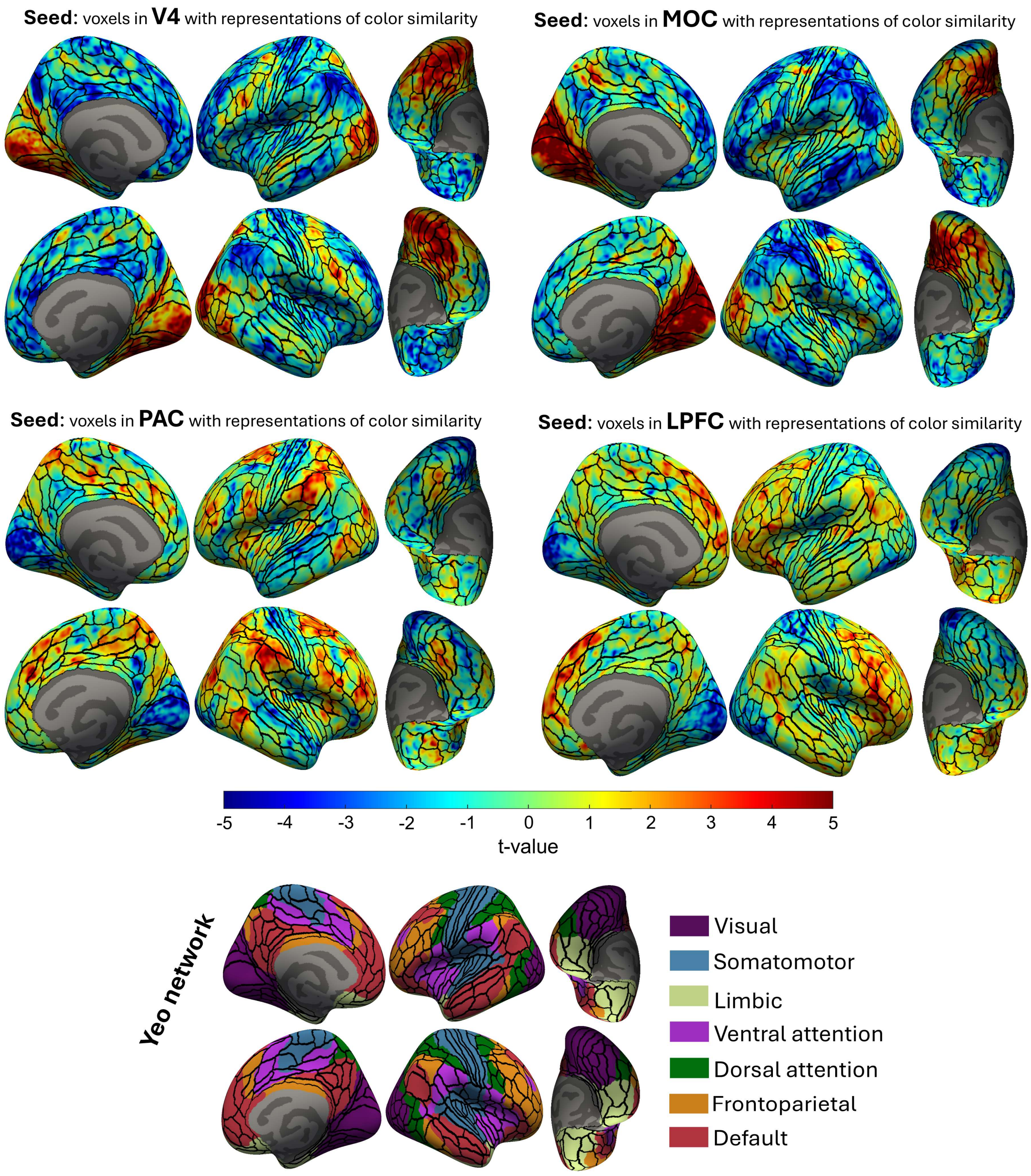


**Supplementary Figure 8. Group-level (N = 8) functional connectivity (FC) map showing the connectivity between voxels with representations of color similarity within an ROI (seed) and the whole brain.**

The FC maps are shown for V4, MOC, PAC, and LPFC. fMRI data were acquired while participants watched the first 20 minutes of the movie Interstellar (the first cohort watched the first 15 minutes). The data underwent our preprocessing steps (see section 4.1.3.1) and were further band-pass filtered (0.01-0.1 Hz) before being used to compute the FC maps. The seed in each ROI included voxels from both left and right hemispheres whose neural RDMs (derived from the RS paradigm) matched the behavioral RDMs with a $p < 0.05$ in a Mantel's test applied to each voxel separately. This threshold ($p < 0.05$; one-tailed as the negative tail is not theoretically expected) was chosen to identify a sufficiently representative subset of voxels; more stringent (e.g., FDR) corrections were overly restrictive, potentially leaving no voxels surviving for the connectivity analysis within an ROI. The mean activity of these voxels during the movie watching was used as the seed regressor in a design matrix that also included the whole-brain mean activity (global mean) and band-pass-filtered head-motion parameters and their derivatives as nuisance regressors. An ordinary least squares regression was used to estimate FCs (beta values). A t-test was then applied to individual-level FC maps to obtain group-level maps. The color bar indicates the t-value from this group-level t-test. The individual participants' FC maps were projected onto an average brain (FreeSurfer's fsaverage), with 5 iterations of surface smoothing applied both before and after projection as implemented in FreeSurfer's mri_surf2surf tool, prior to performing the group-level t-test. All maps are shown unthresholded and are intended for qualitative visualization. The Yeo network parcellation (Yeo et al., 2011) is also shown for comparison of Glasser's atlas (Glasser et al., 2016) regions and the FC maps above with standard networks.

Qualitatively, voxels in V4 with representations of color similarity showed functional connectivity with some areas within MOC (notably around the POS, particularly DVT), within the PAC (around the intraparietal sulcus), and posterior parts of the LPFC. Voxels in the MOC notably showed connectivity with the ventral visual stream, particularly anterior to V4. As for the PAC and LPFC, these are large ROIs (Supplementary Figure 3B) known to comprise various functional subregions. Therefore, the resulting connectivity maps could reflect the functional network within each PAC and LPFC to which the color-representing voxels likely belong. The connectivity map of the PAC showed clusters around the intraparietal sulcus, and the LPFC connectivity map exhibited hotspots in its posterior and middle parts.

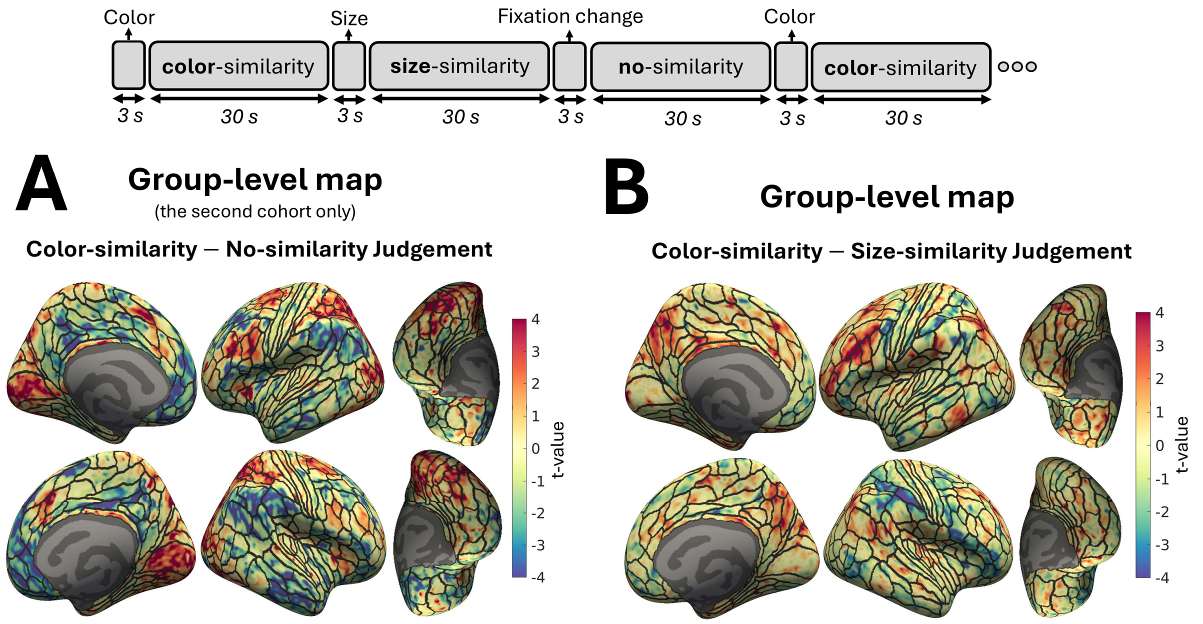


**Supplementary Figure 9. Exploratory maps related to the color-similarity judgment task.** Participants performed 30-second blocks of the color-similarity judgment task, interleaved with two other control blocks of the same (30-second) duration inside the fMRI scanner. In the color-similarity judgment task, participants performed a 2-AFC task, choosing which of two presented colors was more similar to a displayed target color in each trial (Supplementary Figure 1). The colors were displayed as filled circles with slightly varying sizes (a few pixels difference in radius). In one control block, participants judged similarity based on circle size rather than color, selecting the circle closer in size to the target. In the other control block, participants made no similarity judgments. They just viewed the trio of color circles and pressed a key as soon as they noticed a change in the size of the fixation cross displayed at the center of the screen (the fixation cross was also present in the other blocks). After each key press, a new trial (i.e., a new trio of color circles) appeared, just as it would after the 2-AFC response in the other two tasks, color-similarity and size-similarity. Before each block started, participants were informed of the upcoming block type (color-similarity, size-similarity, or no-similarity judgment) by a text cue displayed on the screen for 3 seconds. Participants completed five blocks of each task type, interleaved, within a run, and performed four runs in total, each conducted in a different session using a different color set each time (see Figure 1B). In the first cohort, instead of the no-similarity control block, there was a block in which participants made similarity judgments based on the luminance of the colors (see the non-isoluminant color set in Supplementary Figure 2). They performed one or two runs across their participation sessions, with each run including the color-similarity block twelve times and each of the other blocks six times. See section 4.1.2.2 for a more comprehensive description.

In panels A and B, group-level contrast maps are shown for **(A)** the color-similarity versus no-similarity (N = 4) and **(B)** the color-similarity versus size-similarity judgment task (N = 8). The maps were generated by applying a group-level t-test to the corresponding beta contrast values from each participant (design matrix included one regressor per block type). All maps are displayed unthresholded and are intended for descriptive visualization. The individual participants' beta estimates were projected onto an average brain (FreeSurfer's fsaverage), with 5 iterations of surface smoothing applied both before and after projection as implemented in FreeSurfer's mri_surf2surf tool, prior to performing the group-level t-test. AFNI's afni_proc.py tool was used to obtain beta estimates for each individual, incorporating the same preprocessing steps and nuisance regressors described in section 4.1.3.1.

Qualitatively, the contrast map comparing the color-similarity judgment task to the no-similarity judgment control task (i.e., viewing stimuli without making similarity judgments) showed activity overlapping with regions that appear to contain representations of perceptual similarity (Figure 4B). Specifically, effects were observed in most visual areas, as well as in regions around the intraparietal sulcus in the PAC and in some regions of the posterior LPFC. The contrast map with the control task of judging size-similarity (i.e., a more controlled comparison that involved similarity judgments based on a non-color criterion) showed more limited activity in visual areas. Notably, activity was observed around the intraparietal sulcus, in particular its medial and posterior parts, as well as in the posterior and middle parts of the LPFC, primarily in the left hemisphere. Overall, these results suggested that some of the regions with representations of perceptual similarity may also be engaged during the active task of judging color similarity.

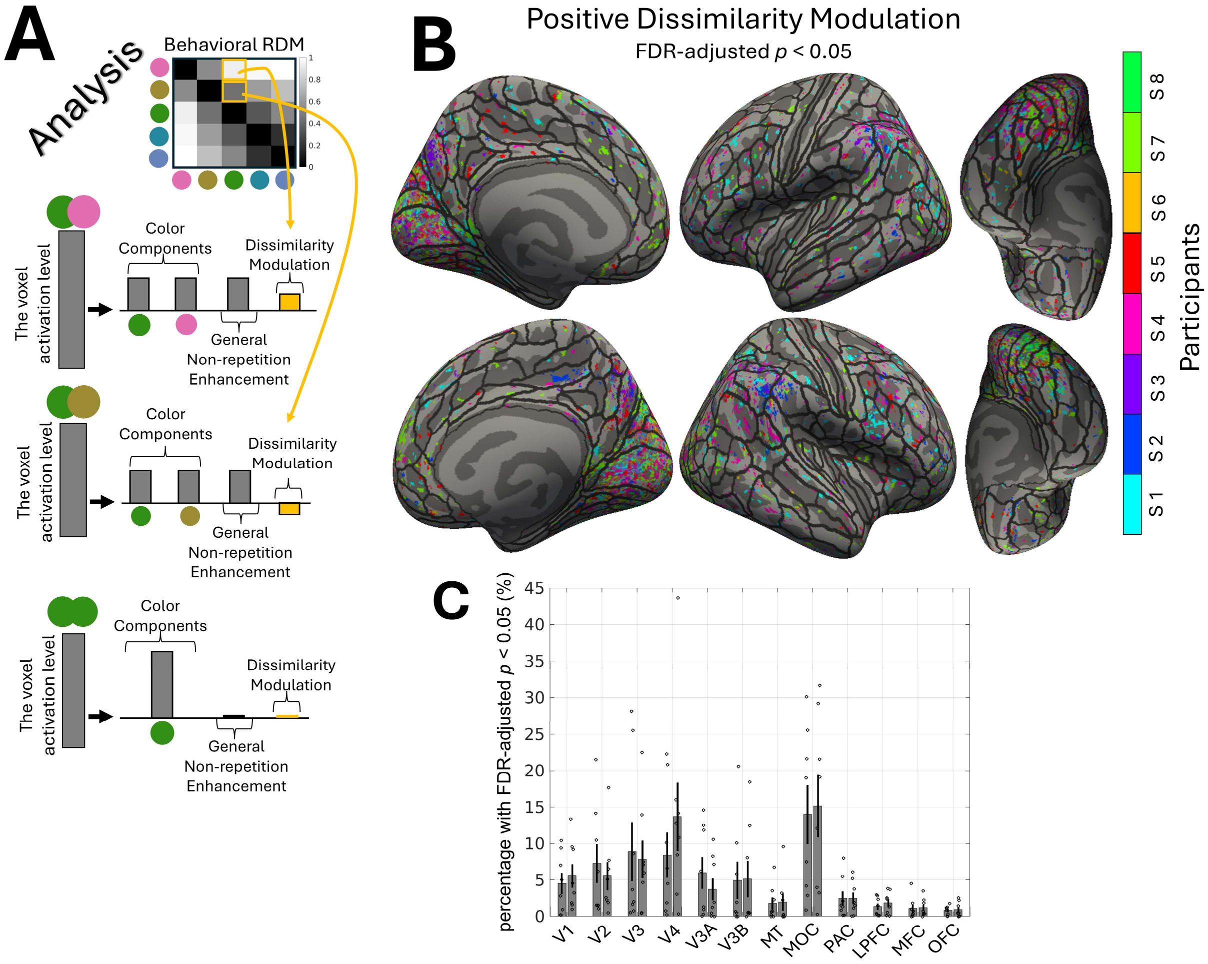


**Supplementary Figure 10. pre-registered univariate analysis of the RS-fMRI data.**

We had pre-registered a 'univariate' analysis that included a regressor in the design matrix to directly reflect voxels with RS profiles aligned with behavioral RDMs. While this analysis is conceptually sound, we found it to be inefficient for the reasons outlined below. Therefore, we did not adopt it as our primary analysis in this paper; instead, we used the RS-MVPA approach, which we found to be more suitable and efficient. First, we realized that some regressors have worrisome multicollinearity, which could make estimates unstable and increase the risk of missing or misattributing effects. Second, we found it inefficient to conduct a group-level analysis following the univariate analysis, given the spatial sparsity and inter-individual variability of the relevant voxels, especially in higher-order areas. These issues reduce reliability and may lead to overlooking relevant regions. In contrast, in the main analysis (Figure 2), the design matrix used to derive the beta estimates (betas used in Figure 2A,1) did not suffer from multicollinearity, and the MVPA approach addressed spatial sparsity and inter-individual variability through an ROI-level analysis.

**(A)** This panel shows the schematic of the analysis approach. Color regressors, one for each color in the color set, were modeled and were activate when the stimulus block contained the corresponding color.

Each color regressor was set to have twice the amplitude when the stimulus block contained only that color to account for its longer (i.e., doubled) total appearance compared to when the color alternated with another color in color-pair blocks. Given the RS principle, a voxel's fMRI activity is expected to be higher when the stimulus block includes two alternating colors than when each color is presented alone. Therefore, one regressor, termed general non-repetition enhancement (GNRE), was included to model this factor, being active in color-pair blocks. Additionally, the RS principle suggests that color pairs with more dissimilar representations within a voxel would result in less suppression, and thus higher fMRI activity, than color pairs with less dissimilar representations (Barron et al., 2016). Therefore, a regressor that modulates the GNRE was included, with its modulation reflecting the behavioral RDM. Consequently, this regressor, termed dissimilarity modulation (DM), indicates how well color representations in each voxel were aligned with the actual perceptual color similarity structures. In summary, the color regressors capture baseline color activation, and the combination of the GNRE and DM regressors identifies voxels with representations of color similarity. A design matrix including these regressors was separately constructed for each color set. The resulting matrices were then combined into a block-diagonal design matrix to form the full model. The GNRE and DM effects were then tested using contrasts that averaged the corresponding regressors across color sets. AFNI's afni_proc.py tool was used to obtain the parameter estimates, incorporating the same preprocessing steps and nuisance regressors described in section 4.1.3.1. Note that while AFNI's 3dREMLfit function (with pre-whitening) tends to yield more conservative individual-level statistics, we employed the default afni_proc.py GLM (no pre-whitening) to prioritize sensitivity, particularly because the non-negligible multicollinearity among regressors already reduces statistical power. **(B)** Voxels with a GNRE t-value greater than 1 were pre-selected, and among them, voxels showing a positive DM at FDR-adjusted $p<0.05$ were identified (one-tailed, because negative DM is not theoretically expected). The results from all participants were superimposed onto an average brain (no smoothing was performed when projecting onto fsaverage), with a different color used to represent each participant. As discussed in panel A, voxels responsive to color should show a positive GNRE; thus, pre-selecting voxels with a GNRE t-value above 1 constrained the FDR correction to potentially relevant voxels with at least some evidence of color responsiveness in the first place. **(C)** In each pre-registered ROI (Supplementary Figure 3B), the percentage of voxels with positive DM (FDR-adjusted $p<0.05$) was calculated. The bar shows the mean percentage across participants, the error bar indicates the standard error, and each dot represents an individual participant. The left and right bars correspond to the left and right hemispheres, respectively.

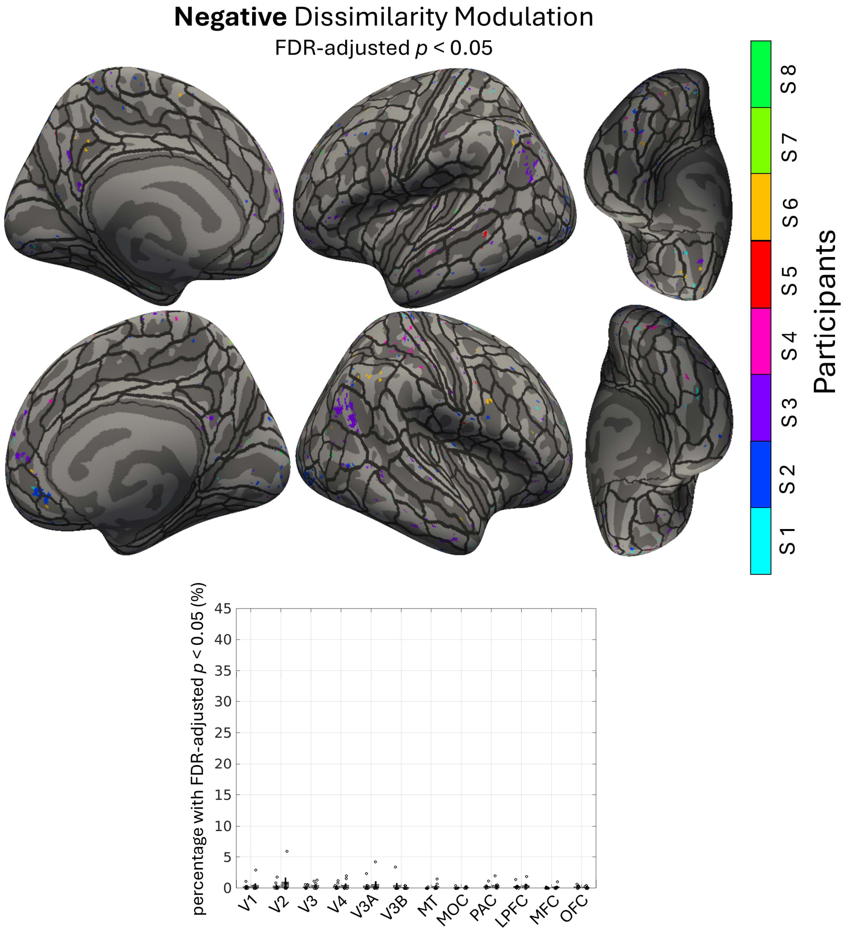


**Supplementary Figure 11. Same as Supplementary Figure 10, but showing the voxels with 'negative' DM (FDR-adjusted $p < 0.05$).**

Although theoretically not expected, we tested for negative DMs (FDR-adjusted $p < 0.05$; one-tailed, focusing on the negative tail), similar to Supplementary Figure 10B, as a sanity check**.** Negative DM reflects representations that are inversely related to perceptual color similarity. The number of voxels with negative DM was negligible and substantially fewer than the number of voxels with positive DM (see Supplementary Figure 10B and 10C). This provides a useful sanity check, as it would be counterintuitive to observe widespread voxels with reversed representations of similarity across participants.

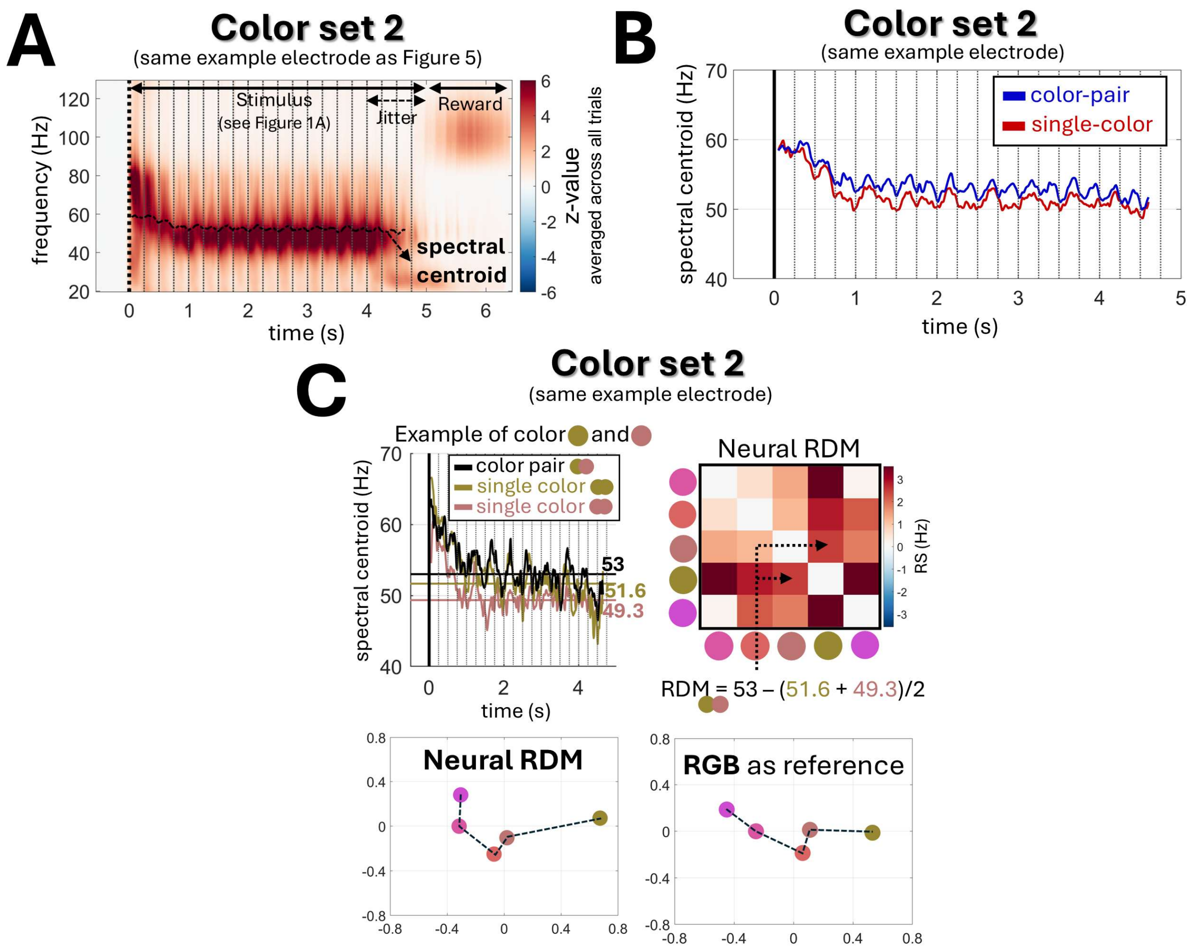


**Supplementary Figure 12. Same as Figure 5A-C, but for color set 2.**

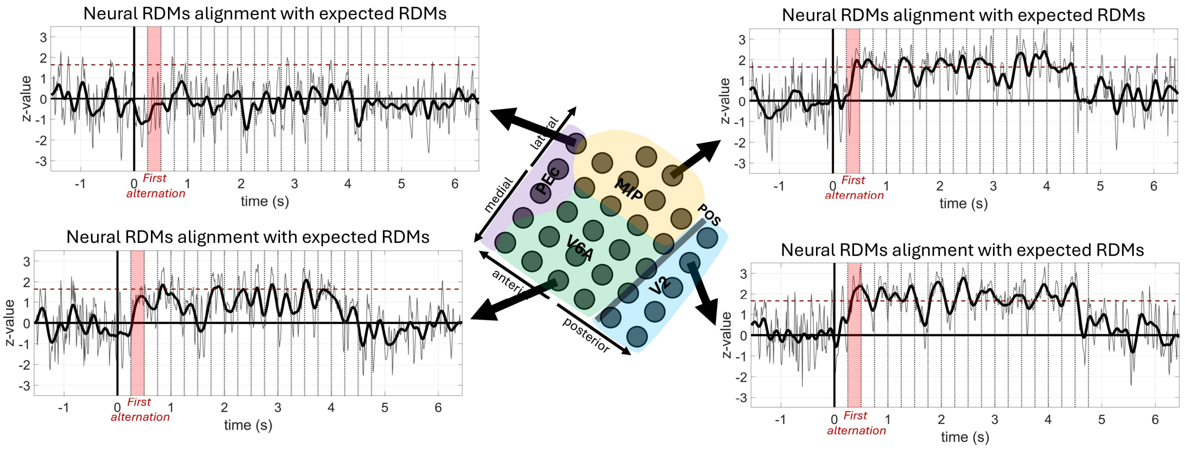


**Supplementary Figure 13. Alignment between extracted neural RDMs and expected RDMs at each time point.**

Neural RDMs were derived from spectral centroid values at each time point (similar to Figure 5C, but using the value at each time point). Subsequently, their alignment with expected RDMs was assessed using Mantel's test, similar to Figure 6C. Each panel shows an example electrode. The y-axis shows Mantel's test z-values and the x-axis shows time. Time 0 s indicates the stimulus onset, and dotted black lines denote color-flickering onsets (every 250 ms). The thin gray curve shows the z-values at each time point, and the thicker black curve shows the smoothed version of the gray curve to reduce noise-related fluctuations and facilitate better visualization of temporal patterns. The broken horizontal red line indicates the z-value of 1.65, corresponding approximately to a *p*-value of 0.05.

As shown here, the RDM derived from the first color alternation already showed apparent alignment with the expected RDMs. This indicates that RS induced by presenting the first color (0-250 ms) followed by the second color (250-500 ms) was already sufficient to measure meaningful representational dissimilarity. The apparent effect could be detected throughout the stimulus presentation period that included color alternation (from 250 ms to about 4,500 ms; stimulus duration was jittered between 4 to 5 s across blocks). We note that, as in Figure 6C, the example electrode in PEc did not show any apparent effect. Also, generally, the effects shown in Figure 6C are stronger than those shown here because Figure 6C used the median spectral centroid value across time. Taking the median across time naturally reduces noise, as it incorporates estimates from several color alternations, resulting in a cleaner estimate of the RDMs.

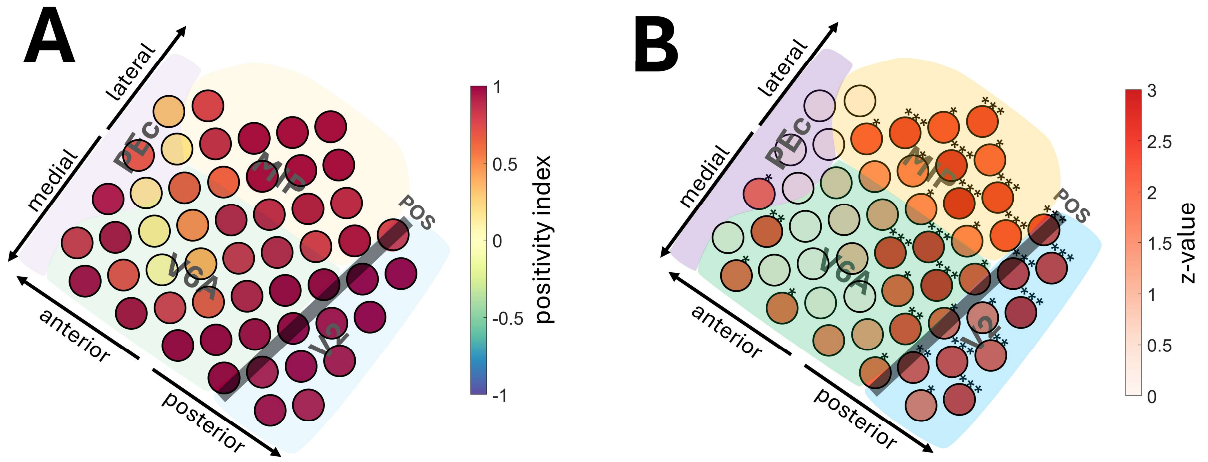


**Supplementary Figure 14. Bipolar analysis.**

Panels A and B are shown in the same manner as Figures 6B and 6C, respectively, but using bipolar electrodes (Mantel's test FDR-adjusted *$p$<0.05, **$p$<0.01, ***$p$<0.005). Signals from adjacent electrode pairs were subtracted, once for vertically adjacent pairs and once for horizontally adjacent pairs, to make a total of 52 bipolar electrodes. The locations of bipolar electrodes were defined as the midpoint between electrode pairs, while the anatomical landmarks (V2, V6A, MIP, PEc) were kept identical as in Figure 6.

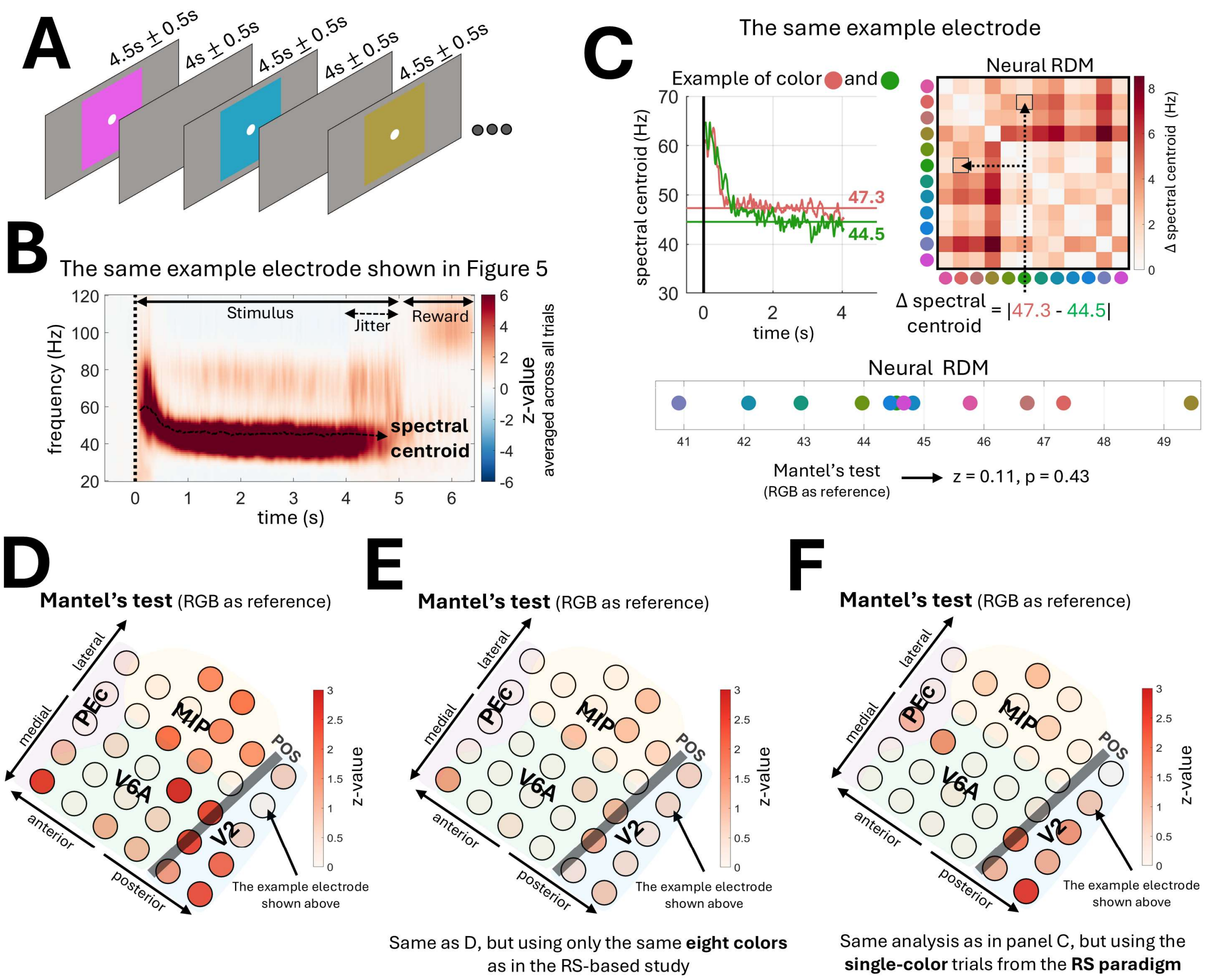


**Supplementary Figure 15. Comparison with a non-RS approach based on evoked responses to individual colors.**

**(A)** In a separate experiment, the NHP viewed each color individually for 4.5 ± 0.5 s, presented as a solid color square and received a reward for maintaining central fixation. Unlike the RS paradigm, this experiment did not use checkerboard color flickering and included 12 isoluminant colors (Figure 1B), rather than only the eight colors from sets 1 and 2. **(B)** This panel shows the time-frequency response, z-normalized relative to pre-stimulus baseline and then averaged across all trials for an exemplar electrode in V2 (the same electrode as in Figure 5). **(C)** The median spectral centroid value during the steady period (1.5 s after stimulus onset) corresponding to each color was used to construct a 12 × 12 RDM. Each cell in the RDM was computed as the absolute difference between the median spectral centroid values of a pair of colors. This panel shows the RDM for the same exemplar electrode, visualized in 1D space. Note that in this approach, the representation at a single electrode is inherently one-dimensional because of how it is constructed and, unlike in the RS-based approach, cannot be multidimensional. For the exemplar electrode shown, although the resulting representation shows a discernible pattern, with lower spectral centroid values for blue/greenish colors and higher values for orange/pinkish colors, its alignment with an RGB-based reference representation of color similarity is low (Mantel's test: z = 0.11, p = 0.43). **(D)** This panel shows the alignment between the RDM derived from each electrode (as shown in panel C) and the RGB-based reference representation of color similarity. Although some electrodes showed an apparent effect (uncorrected $p < 0.05$), no electrode passed the FDR-adjusted threshold of $p < 0.05$ in the Mantel's test. **(E)** Same as panel D, but only eight colors were used (an 8 × 8 RDM), corresponding to the same colors used in the RS-based approach. Limited effects were observed, with none passing the FDR-adjusted threshold of $p < 0.05$. **(F)** The same analysis was applied (i.e., the RDM was constructed using the same procedure as shown in panel C) but using single-color blocks from the RS paradigm (i.e., blocks in which a single color was presented). No electrode passed the FDR-adjusted threshold of $p < 0.05$.

This figure shows that a univariate, non-RS method that constructs representations from single-color-evoked responses may provide insufficient resolution to detect representations of color similarity. In other words, this highlights the power of RS in recovering high-resolution representations that are not accessible with approaches such as the one shown here.

Notably, this non-RS method showed stronger results when a multivariate approach was applied based on response patterns across electrodes rather than considering each electrode individually. The responses (i.e., the median spectral centroid values) across electrodes were treated as a vector, yielding one response vector per color. The distances (1 - correlation) between these vectors were then used to construct an RDM. When all 12 colors were used, the alignment with the RGB-based reference representation of color similarity was significant (Mantel's test: z = 3.1, p = 0.003). However, when the same eight colors as in the RS-based approach were considered, the alignment did not reach significance (Mantel's test: z = 0.95, p = 0.18). Therefore, even with the multivariate analysis, this non-RS method was less sensitive than the RS-based approach. Similarly, when this multivariate analysis was applied using single-color trials from the RS paradigm, the alignment was trend-level and did not reach significance (Mantel's test: z = 1.1, p = 0.14). We note that using alternative distance measures other than correlation (e.g., Euclidean distance) did not yield significant results, even in the case of 12 colors.

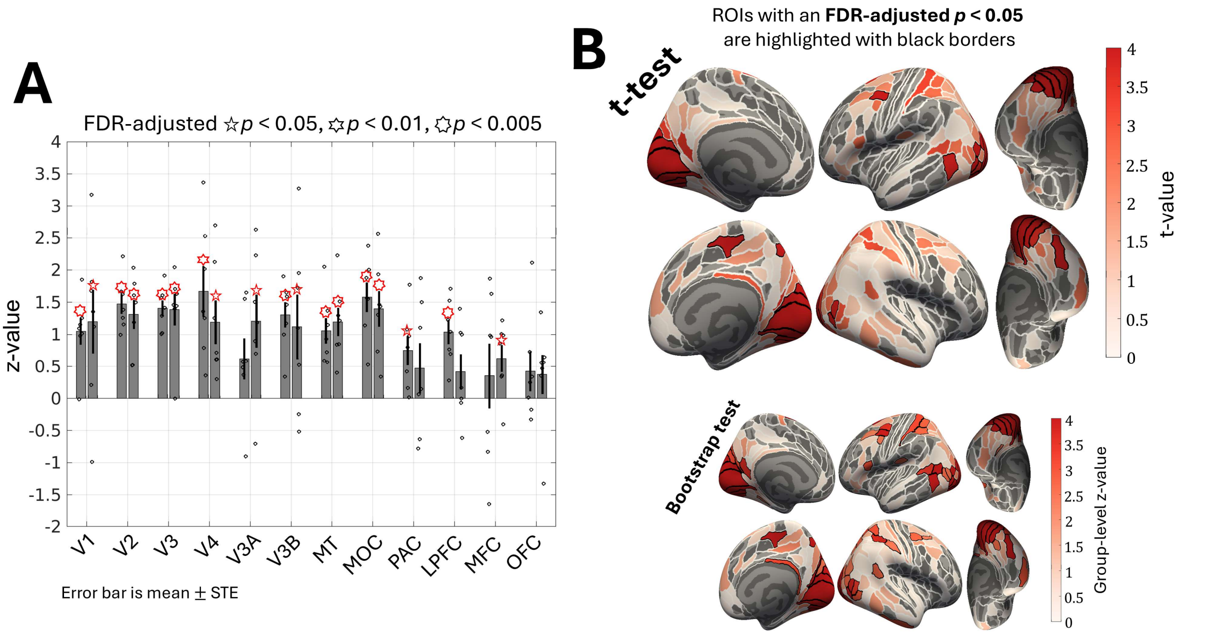


**Supplementary Figure 16. Same as Figure 4, but including only isoluminant color sets in the analysis (based on three participants with set 2 from the first cohort and the entire second cohort; see section 4.1.2.1).**

In panel B, atlas results based on a bootstrap test are additionally shown. Bootstrap test provides higher sensitivity at the group-level than a t-test, although this does not necessarily imply a better performance.

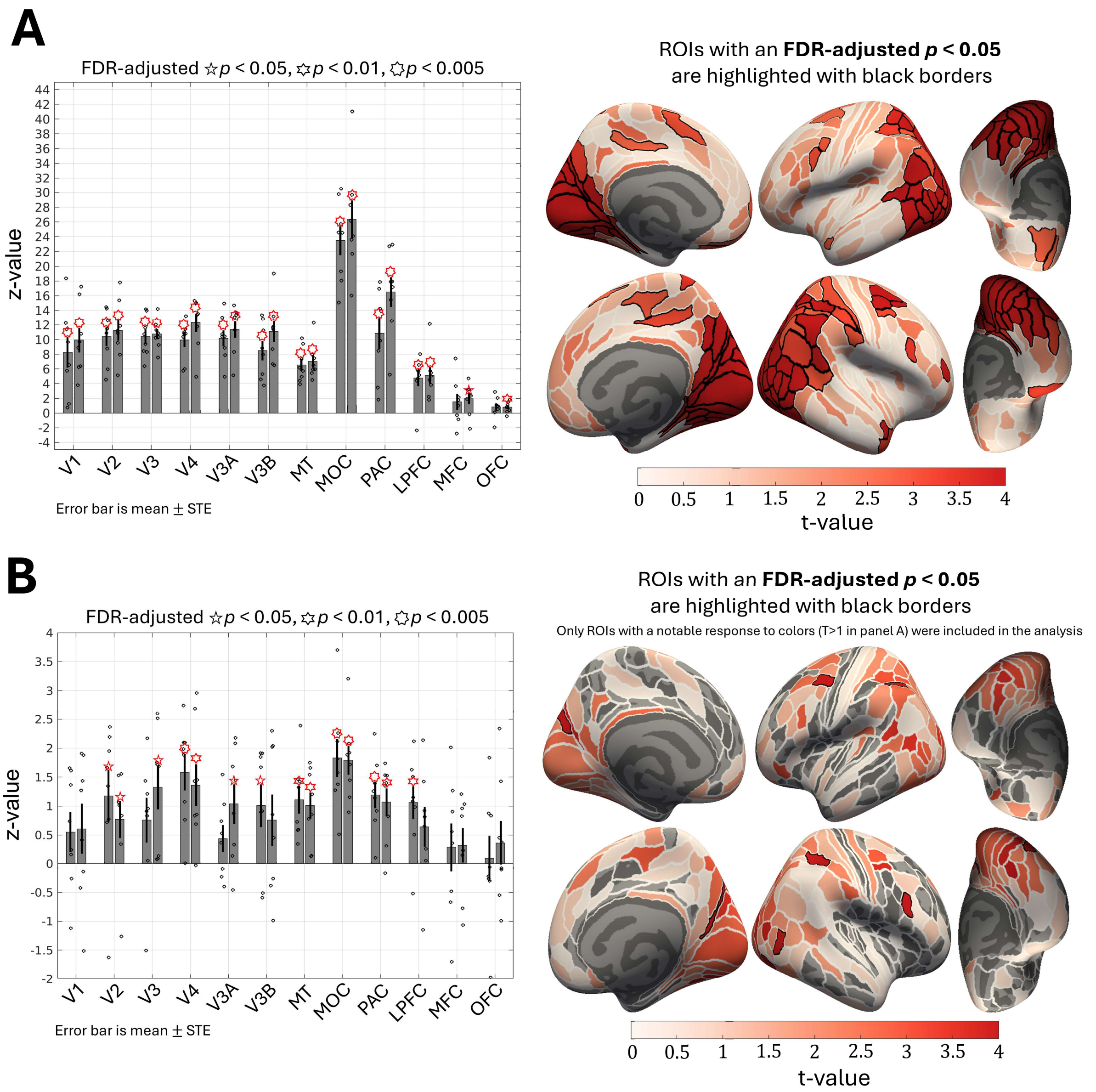


**Supplementary Figure 17. Panels A and B correspond to Figures 3 and 4, respectively, but show the results when a typical GLS model-fitting was used to derive beta estimates (see section 4.1.3.1).**

The plot descriptions are the same as those in Figures 3 and 4.

**Supplementary video.** Examples of stimuli presented in the RS paradigm, including colors from color set 3 (see Figure 1B). Note that the example colors shown in the video may not appear isoluminant due to variations in monitor settings. However, the colors presented in the experiment were physically isoluminant.
https://drive.google.com/file/d/1Phdgmbaot167dj5JjVVI5f96ie_PK7RQ/view?usp=sharing
